# H2A.Z-H2B.V nucleosomes form a specialised scaffold for transcription initiation in *Trypanosoma brucei*

**DOI:** 10.64898/2026.09.18.752700

**Authors:** Gauri Deák Ünal, Christos Spanos, Martin R. Singleton, Marcus D. Wilson

## Abstract

Regions of transcription initiation are often enriched in the histone variant H2A.Z. This is especially true in the divergent eukaryote *Trypanosoma brucei*, where broad domains of H2A.Z and a kinetoplastid-specific variant H2B.V define a limited number of polycistronic transcription start sites. To explore the chromatin underpinning this unusual transcription, we reconstituted H2A.Z-H2B.V nucleosomes *in vitro* and found that they are inherently unstable, form open chromatin structures, and serve as a direct interaction scaffold for gene regulators, in line with their role in transcription activation. A single particle cryo-EM structure reveals why H2A.Z-H2B.V form obligate dimers, pack differently within nucleosomes, and exhibit altered binding to DNA. We identified unique features of H2A.Z and H2B.V that result in differential activity of chromatin modifying enzymes, including the H2A.Z C-terminal tail and an atypical acidic patch. An affinity purification mass spectrometry screening approach revealed that H2A.Z-H2B.V nucleosomes have a distinct protein interaction profile, suggesting that interactions with chromatin factors are directly tuned for transcription control. Overall, this work reveals how trypanosome-specific histone features both inherently alter the biomechanical properties of nucleosomes as well as reshape chromatin binding and modification patterns at transcription start sites.

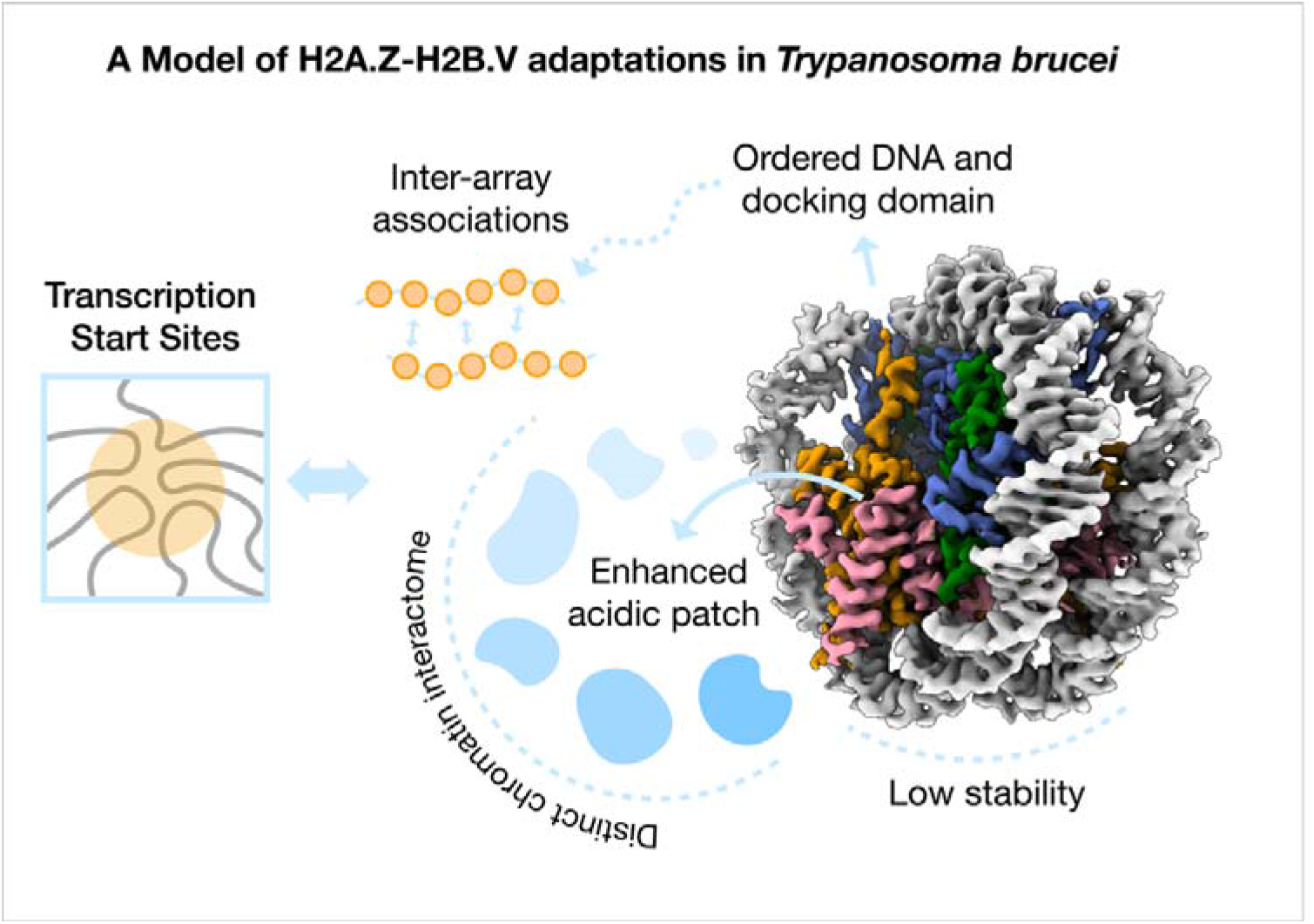
Graphical Abstract.

## Introduction

Chromatin structure plays a fundamental role in regulating genome function. By packaging DNA into nucleosomes, chromatin restricts access to genetic material and acts as a landing platform to influence DNA related processes. As a result, modulation of nucleosome stability and dynamics is a central mechanism by which eukaryotic cells regulate these processes. Histone variants contribute substantially to this regulation by altering nucleosome structure and function, generating chromatin states with distinct physical and biochemical properties (1,2).

Among histone variants, H2A.Z is one of the most evolutionarily conserved and extensively studied (3,4). H2A.Z has been implicated in diverse and sometimes apparently contradictory functions, depending on its specific genomic context. This includes roles in transcriptional activation and repression, RNA polymerase II (RNAPII) pausing and elongation, DNA repair, and higher-order chromatin organisation (4). In many eukaryotes, incorporation of H2A.Z alters nucleosome stability and positioning, producing nucleosomes that are generally more dynamic and accessible than their canonical counterparts (5). H2A.Z occupancy frequently correlates with transcriptionally active promoter regions and paused RNAPII, suggesting an important role in regulating the balance between chromatin accessibility and transcriptional control (6–8). Notably, subtle sequence differences between H2A.Z orthologues and isoforms can profoundly influence nucleosome stability, genomic localisation, and protein interaction networks (9–11). As such, studying H2A.Z sequence variation and function in different evolutionary contexts can reveal important aspects of H2A.Z function as well as the general principles of histone variant evolution.

*Trypanosoma brucei*, the causative agent of African trypanosomiasis, exhibits a highly unusual mode of gene expression that differs from well-studied eukaryotes. Protein-coding genes are arranged in large polycistronic transcription units that are constitutively transcribed, while regulation of gene expression occurs predominantly through post-transcriptional mechanisms (12,13). Transcription initiates at a limited number of transcription start site regions (TSS), which are defined by specialised chromatin rather than sequence-specific transcription factors (14–17). Notably, trypanosomes are uniquely reliant on histone variants to demarcate different areas of the genome (14). Indeed, sequence divergence within trypanosome histones and histone variants can directly reshape nucleosome structure and stability, thereby encoding biological function at the level of chromatin architecture (18–20).

A hallmark of trypanosomatid TSS is the enrichment of the histone variants H2A.Z and a H2B variant termed H2B.V. H2A.Z and H2B.V occupy broad domains spanning ∼10 kb and demarcate sites of transcription initiation (14,15,21,22). The two variants function as a pair in *T. brucei* and are essential in several trypanosomatids (22–24). Perturbing trypanosome H2A.Z alters TSS positioning, reduces mRNA levels and changes chromatin structure (15,25,26). Rather than being restricted to narrow promoter regions, H2A.Z is present throughout these extended transcription start domains, which cluster in three dimensions (27), suggesting that H2A.Z-H2B.V contributes to a distinct chromatin environment associated with constitutive transcription initiation. H2A.Z-H2B.V-containing chromatin is also more accessible than surrounding genomic regions (15) and is enriched in numerous chromatin factors involved in writing, reading, and erasing histone modifications (28), underscoring the central role of these variants in trypanosome biology.

The remarkable conservation of H2A.Z across eukaryotic evolution suggests that its ability to modulate chromatin structure represents a fundamental feature of genome regulation (4,29). Structural and biophysical studies in fungi, plants and mammals have defined how H2A.Z tunes nucleosome stability, DNA accessibility and effector binding (5,11,30–34). Whereas H2A.Z is conserved across eukaryotes, its trypanosome partner H2B.V is restricted to kinetoplastids and forms a variant pair that is rarely found in eukaryotes (14,23). Whether the structural and functional properties of H2A.Z-H2B.V nucleosomes reflect conserved mechanisms of H2A.Z action or have evolved lineage-specific adaptations remains unknown. Resolving this question is important both for understanding transcriptional regulation in trypanosomes and for establishing how evolutionary diversification of histone variants can generate novel chromatin states.

Here, we sought to define the molecular basis of H2A.Z-H2B.V function using complementary biochemical, structural, and proteomic approaches. We reconstituted nucleosomes containing *T. brucei* H2A.Z and H2B.V and demonstrated that these variants form an obligate heterodimer *in vitro*. The structure of the H2A.Z-H2B.V nucleosome reveals how divergent sequence features encode altered chromatin properties, including reduced nucleosome stability, altered properties of nucleosome arrays, and different engagement by chromatin enzymes and effector proteins. We found that H2A.Z-H2B.V nucleosomes directly interact with chromatin factors that are commonly enriched at transcription start regions, including bromodomain proteins and a SIN3-like histone deacetylase complex. Together, our findings reveal how evolutionary diversification of H2A.Z and H2B.V generates a specialised chromatin state that defines TSS in *T. brucei* and provides broader insights into the mechanisms by which histone variants shape chromatin function.

## Materials and Methods

### Protein Sequence Alignments

Kinetoplastid protein sequences were retrieved from TriTrypDB (35) and human sequences were retrieved from UniProt (36). Sequence-based alignments were performed with MAFFT and visualised in JalView (37). Pairwise percentage sequence identity values were obtained using MUSCLE (38) and visualised using the Bioconductor package ComplexHeatmap (39) and the viridis colour scheme. Dendrograms were generated using hierarchical clustering by Euclidean distance.

### Protein Structure Prediction and Alignments

Protein structure predictions were retrieved from the AlphaFold2 database (40). The predicted structure of the HDAC1 complex member Q57TZ5 was used to query the AlphaFold and PDB databases for potential structural homologs using Foldseek (41). Structure-based alignments of Q57TZ5 and SIN3A proteins were performed with TM-Align using the RCSB server (42).

### Plasmid Constructs

Information about all plasmid constructs (including their original source, backbone, encoded protein/DNA, mutations, and PCR primers) is provided in Supplementary Data File 1. New plasmids made for this study were prepared by cloning a gBlock Gene Fragment (Integrated DNA Technologies) into a plasmid backbone using the NEBuilder® HiFi DNA Assembly Cloning Kit (New England Biolabs). Point mutations were introduced by site-directed mutagenesis.

### Protein Purification

#### Histones

Histones were purified from inclusion bodies as described previously (19,20,43). Briefly, histones were expressed in *Escherichia coli* BL21(DE3)RIL cells for 3-4 h at 37°C. Cells were lysed and inclusion bodies solubilised in guanidine hydrochloride based unfolding buffer. The supernatant was dialysed into Ion Exchange Buffer A (15 mM Tris pH 7.5, 7 M urea, 100 mM NaCl, 1 mM EDTA, and 5 mM 2-Mercaptoethanol), loaded onto a HiTrap SP XL column (Cytiva), and eluted with a 0-80% gradient of Ion Exchange Buffer B (15 mM Tris pH 7.5, 7 M urea, 1 M NaCl, 1 mM EDTA, and 5 mM 2-Mercaptoethanol). Fractions were analysed by SDS-PAGE and the purest fractions were dialysed into 1 mM acetic acid, lyophilized, and stored at −20°C.

Unless stated otherwise, N-terminal His_6_-TEV tags on *Tb* H2A.Z histone constructs were cleaved as follows. Lyophilized histones were resuspended in Histone Cleavage Buffer (20 mM Tris pH 7.5, 1M urea, 100 mM NaCl, 4 mM sodium citrate, and 2 mM 2-Mercaptoethanol) and incubated with TEV protease at 4°C overnight. The reaction was adjusted to match Histone IMAC Buffer A (20 mM Tris pH 7.5, 5M urea, 500 mM NaCl, 25 mM imidazole, and 2mM 2-Mercaptoethanol) and flowed through a nickel-charged HiTrap IMAC HP column (Cytiva). The flow-through containing cleaved histones was collected and remaining protein was eluted with Histone IMAC Buffer B (20 mM Tris pH 7.5, 5M urea, 500 mM NaCl, 300 mM imidazole, and 2mM 2-Mercaptoethanol). The reaction products were checked by SDS PAGE and stained with a Colloidal Coomassie Stain (10% tartaric acid, 2% ethanol, 1% alpha-cyclodextrin, 0.3% hydroxyethyl cellulose ∼90,000 Da, 0.0015% Coomassie G250 dye w/v). Pooled fractions were then dialysed into 1mM acetic acid, and lyophilized.

#### His_6_-MBP-TEV (HMT)-DOT1B

*T. brucei* DOT1B (UniProt ID: Q4GZF2, TriTrypDB ID: Tb927.1.570) was purified with an N-terminal His_6_-MBP-TEV tag as described previously (20). Briefly, HMT-DOT1B was expressed in *E. coli* BL21(DE3)RIL cells, lysed and the soluble fraction was loaded on a nickel-charged HiTrap^TM^ HP IMAC column (Cytiva) and eluted with a gradient of imidazole. The protein was further purified by ion-exchange chromatography and size exclusion chromatography in 15 mM HEPES pH 7.5, 150 mM NaCl, 5% glycerol (v/v), and 2 mM DTT. Fractions were analysed by SDS-PAGE. The purest fractions were then spin-concentrated, and stored at −80°C.

#### KKT23

*T. brucei* KKT23 (UniProt ID: Q38AU6, TriTrypDB ID: Tb927.10.6600) was purified as described previously (44), by TALON affinity chromatography prior to tag removal by TEV protease and further purification using ion exchange and size exclusion chromatography. The final buffer comprised 50 mM Tris pH 8, 50 mM NaCl, 5% glycerol (v/v), 1 mM DTT, 0.1mM EDTA, and 10mM sodium butyrate.

### Histone Fluorescent Labelling

*Tb* H2A K120C was labelled with Alexa Fluor^TM^ 647 C_2_ maleimide (Cat#A20347, Thermo Fisher Scientific, hereafter “Alexa647”) or Oregon Green 488 maleimide (Cat#715, AAT Bioquest®, hereafter “OGG488”). *Tb* H2A.Z K169C was labelled with OGG488 maleimide. Labelling was performed as described previously(45,46), with some modifications. Lyophilized histones were resuspended in Maleimide Reaction Buffer (15 mM HEPES pH 7.0, 7 M guanidine-HCl, 25 mM NaCl, and 0.5 mM TCEP). Each histone (200 μM) was incubated with a fluorescent dye at a 2:1 molar ratio overnight at 4°C in a 1 mL reaction volume. The histones were then combined with H2B or H2B.V in Maleimide Reaction Buffer, refolded into dimers, and purified by size exclusion chromatography (see section Preparation of Histone Dimers, Tetramers, and Octamers). The degree of labelling (DOL) achieved was 33-45% and was calculated as:

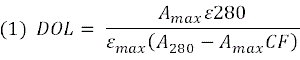

Where:

*A_max_* = histone absorbance measured at the absorbance maximum of the fluorescent dye

A_280_ = histone absorbance measured at 280 nm

*ε_max_* = molar extinction coefficient of the fluorescent dye at its absorbance maximum

*ε_280_* = molar extinction coefficient of the histone at 280 nm

*CF* = correction factor of the fluorescent dye at 280 nm

### Preparation of Histone Dimers, Tetramers, and Octamers

Lyophilized histones were resuspended in Unfolding Buffer (20 mM Tris pH 7.5, 7 M guanidine hydrochloride, and 10 mM DTT) for 30 min at room temperature and combined at different molar ratios to form dimers (1:1 of H2A:H2B), tetramers (1:1 of H3:H4), or octamers (1.2:1.2:1:1 of H2A:H2B:H3:H4). A full list of canonical, variant, or hybrid histone complexes that were prepared is available in Supplementary Data File 1. The histone mixture was then diluted to 2 mg.mL^-1^ and dialysed into Refolding Buffer (15 mM Tris pH 7.5, 2M NaCl, 1mM EDTA, 5 mM 2-Mercaptoethanol). For complexes containing *Tb* H2A.Z (or derivatives), 2-Mercaptoethanol in the Unfolding and Refolding Buffers was replaced with 50 mM DTT and 5 mM DTT respectively due to a high propensity for disulphide bond formation. Following dialysis, the refolded complex was concentrated in an appropriate MWCO Amicon spin concentrator (Sigma Aldrich) and loaded on a Superdex200 or Superdex75 size exclusion column (dimers) and eluted in Refolding Buffer. Fractions were analysed by SDS PAGE, the purest fractions were pooled, and the final complex was spin concentrated. Histone octamers were stored at –20°C in 50% Refolding Buffer, 50% glycerol (v/v). Histone dimers and tetramers were stored at 4°C in Refolding Buffer or flash frozen in liquid nitrogen and stored at –80°C.

### Preparation of DNA for Reconstitutions

#### Nucleosome Core Particle (NCP) and Mono-Nucleosome DNA

DNA for wrapping nucleosomes or NCPs was prepared using a PCR-based approach as described previously (19,20). Briefly, four to eight 96-well plates of 100 µL PCR reactions with Pfu polymerase were loaded on a Resource Q anion exchange column (Cytiva) pre-equilibrated with Q-Buffer A (10 mM Tris pH 8.0, 1 mM EDTA). The DNA was eluted with a 25-45% gradient of Q-Buffer B (10 mM Tris pH 8.0, 1 mM EDTA, 2 M NaCl). Fractions were analysed on a 5% polyacrylamide Tris-glycine native gel and the purest fractions were pooled. The obtained DNA was ethanol precipitated, resuspended to 5-10 mg.ml^-1^ in 10 mM Tris pH 8.0, 0.1 mM EDTA, and stored at –20°C. Biotinylated DNA was prepared by using a 5’ biotin-TEG primer with an 18-atom hexa-ethyleneglycol linker (IDT). A list of DNA sequences and PCR primers is available in Supplementary Data File 1.

#### 12×177 bp Nucleosome Array DNA

12 repeats of Widom 601 177 bp nucleosome array DNA was prepared as described previously (20,47), with some amendments. A plasmid encoding the array DNA flanked by EcoRV restriction sites was amplified in NEB® Stable or TOP10 (Invitrogen) *E. coli* cells and isolated using a MaxiPrep Kit (Qiagen). The plasmid was digested with EcoRV (NEB) overnight at 37°C in NEBuffer™ 3 (50 mM Tris pH 7.9, 100 mM NaCl, 10 mM MgCl2, 1 mM DTT). The array DNA fragment was precipitated by addition of 5.2% PEG6000 and 750 mM NaCl and centrifuged for 1h at 4,000 x g. The insoluble fraction containing array DNA was further purified by ethanol precipitation, and resuspended in 10 mM Tris pH 8.0, 0.1 mM EDTA. The quality of each preparation was checked by agarose gel electrophoresis.

#### 147 bp MMTV Competitor DNA

A plasmid with eight copies of 147 bp MMTV DNA (47) flanked by EcoRV restriction sites was amplified, isolated, and digested with EcoRV as described for array DNA above. MMTV fragments were separated from the plasmid backbone by addition of 9.3% PEG6000 and 500 mM NaCl and centrifuged for 1h at 4,000 x g. The supernatant was retained, ethanol precipitated, and resuspended in 10 mM Tris pH 8.0, 0.1 mM EDTA. The quality of each preparation was checked by agarose gel electrophoresis.

### Reconstitution of NCPs and Nucleosomes

NCPs were prepared with Widom 601 145 bp DNA and nucleosomes were prepared with Widom 603 193 bp DNA or biotinylated Widom 601 175 bp DNA. In the case of human and *Tb* canonical NCPs/nucleosomes, a standard approach was used, where 0.3 mg.mL^-1^ DNA was combined with histone octamers at an optimal molar ratio (0.5-1.0 DNA to 1.0 octamer) (19,20,43). In the case of NCPs/nucleosomes containing *Tb* H2A.Z-H2B.V (or derivatives), octamer formation was unsuccessful, so an alternative method combining separate histone dimers, tetramers, and DNA at a 2:1:(0.5-1.0) molar ratio was used (43). Each mixture was subjected to 18h gradient dialysis from 2 M KCl to 200 mM KCl in 15 mM HEPES pH 7.5, 1 mM EDTA, 1 mM DTT at 4°C. The reconstituted complexes were then dialysed into a final Storage Buffer (15 mM HEPES pH7.5, 25 mM NaCl, 1 mM DTT) for 3h at 4°C. The NCPs/nucleosomes were analysed by SDS PAGE (17% polyacrylamide) and native PAGE (5% polyacrylamide, Tris-glycine gels stained for DNA using Promega Diamond^TM^ Nucleic Acid Dye). The final concentration of NCPs/nucleosomes was based on quantification of double-stranded DNA and was measure in ng.µL^-1^ using a NanoDrop.

### Nucleosome Array Reconstitution

Nucleosome arrays were reconstituted in a similar manner as previously described (20,47). The arrays were prepared by combining optimised molar amounts of 12x Widom 601 177 bp DNA (x1), 147 bp MMTV competitor DNA (x6), histone dimers (x48) and tetramers (x24), or histone octamers (x24). To prepare fluorescent Alexa647-labelled nucleosome arrays, fluorescent *Tb* H2AK120C-Alexa647 dimers were added in limiting amounts (x4) to unlabelled dimers (x44) and tetramers (x24). Alternatively, the *Tb* H2AK120C-Alexa647 dimers (4x) were added to tetramers (x2) and different variant and mutant octamers (x22). OGG488-labeled arrays were reconstituted with *Tb* H2AK120C-OGG488 or *Tb* H2A.ZK169C-OGG488 dimers (x48). The components were mixed and incubated on ice for 30 min in 2M KCl, 15 mM HEPES pH 7.5, 1 mM EDTA, 1mM DTT.

Each reaction was subject to gradient dialysis for 18h from 2 M KCl to 200 mM KCl in 15 mM HEPES pH 7.5, 1 mM EDTA, 1 mM DTT at 4°C and then dialysed for 3h into a final Array Storage Buffer (15 mM HEPES pH 7.5, 25 mM NaCl, 1 mM EDTA, 1 mM DTT) at 4°C. As for nucleosomes/NCPs, the concentration of nucleosome arrays was based on quantification of double-stranded DNA and measured in ng.µL^-1^ using a NanoDrop based on quantification of double-stranded DNA.

To remove excess free DNA, each nucleosome array was precipitated with 1 volume of 2x Precipitation Buffer (14 mM MgCl_2_, 15 mM HEPES pH 7.5, 1 mM EDTA, 1 mM DTT) for 15 min on ice and centrifuged for 15 min at 17,000 x g, 4°C. The resulting pellets were resuspended in Array Storage Buffer and further dialysed into Array Storage Buffer overnight at 4°C. 50 ng of input, an equivalent amount of supernatant, and 50 ng of pellet from each precipitation reaction were analysed on a 0.7% (w/v) low EEO agarose 0.25x TBE native gel and post-stained with Diamond^TM^ Nucleic Acid Dye (Promega).

Saturation of nucleosome arrays was checked as described previously(20,48), with some modifications. The 12x Widom 601 177 bp DNA used to reconstitute arrays contains a single AccI restriction site in the core of each Widom 601 repeat that is normally protected by nucleosomes. To confirm this, 15 μL reactions containing 200 ng of arrays and 10 units of AccI (NEB) were incubated for 1h at 26°C in NEB rCutSmart Buffer and quenched with 4.4 μL of Stop Solution (2.7% SDS, 136 mM EDTA, 4.5 mg.mL^-1^ NEB Proteinase K) for 1h at 37°C. 60 ng of each reaction were then analysed on a 1% (w/v) agarose TAE gel stained with SYBR Safe DNA Gel Stain (Invitrogen). The 12x Widom 601 177 bp DNA also contains a single ScaI restriction site in each linker DNA segment, which should be accessible for digestion and generate intact mononucleosome species. 15 μL reactions containing 200 ng of arrays and 10 units of ScaI (NEB) were incubated for 3h at 26°C in NEB rCutSmart Buffer. 60 ng of each reaction analysed by native PAGE (5% polyacrylamide, Tris-glycine gel) and stained with Diamond^TM^ Nucleic Acid Dye (Promega).

### Single Particle Cryo-EM Sample Preparation and Data Collection

(His_6_-TEV-H2A.Z)-H2B.V NCPs reconstituted with Widom 601 145 bp DNA were cross-linked with 0.03% glutaraldehyde (v/v) for 5 min on ice and quenched with 50 mM Tris pH 7.5 and 50 mM ammonium bicarbonate. The NCPs were then buffer exchanged into Cryo-EM Buffer (15 mM Tris pH 7.0, 25 mM NaCl, 1 mM EDTA, and 1 mM DTT) using a 100 kDa MWCO Amicon spin concentrator (Sigma Aldrich). The final sample was diluted to 110 ng.µL^-1^ and 3.5 µL were applied onto holey carbon 300 mesh R2/2 grids (Quantifoil). The grids were pre-treated by fresh carbon evaporation and glow discharged for 60s at 25 mA, 38 Pa in a PELCO easiGlow^TM^ system. The grids were then blotted and vitrified in liquid ethane cooled by liquid nitrogen using a Vitrobot Mark IV system (4°C, 100% humidity, 3.5 s blotting time).

Grid screening and optimisation was performed in-house, using a 200 kV FEI Tecnai F20 microscope. A single clipped grid was then imaged using a TFS Titan Krios microscope operated at 300 kV and equipped with a Gatan K3 (6k x 4k) direct electron detector. 13,861 movies were collected with a pixel size of 0.829 Å.px^-1^, a 2.3 s exposure time, a total electron dose of 49.53 e^-^.Å^-2^, and a target defocus range of –2.27 to –1.25 μm (0.25 steps) at the Electron Bio-Imaging Centre, Diamond Light Source, UK.

### Single Particle Cryo-EM Data Processing

13,861 micrographs were imported into CryoSPARC (49), motion corrected with Patch Motion Correction, and CTF parameters estimated using Patch CTF Estimation. Micrographs with low CTF fit resolution, high ice contamination, and aggregation were discarded and 9,924 micrographs were taken forward. Manual picking (945 particles) was performed to generate 2D class templates for template-based picking (714 particles). A sample of 200 micrographs was used to train a Micrograph Denoiser Model (CryoSPARC implementation) and 3,094,412 particles were picked on denoised micrographs using the prepared 2D class templates and extracted on non-denoised micrographs with a boxsize of 256 px binned by 4. Two rounds of 2D classification with 100 classes were performed (Round 1: 1,200,200 particles retained, Round 2: 1,055,641 particles retained). *Ab initio* reconstruction was used to generate an initial model. Particles were extracted with a box size of 256 px (at 0.829 Å.px^-1^), homogenously refined and subject to 3D classification. Two 3D classes with better resolution of structural features were selected for further refinement (585,210 particles). Particles from these classes were re-extracted with a box size of 304 px (574,920 particles retained) and used to generate a model by homogeneous refinement. Reference-based motion correction was applied and the particles were further separated into two 3D classes. One 3D class was selected (339,596 particles) and particle orientations were rebalanced retaining the top 90% particles in the most over-represented bins based on orientation alignment to reduce orientation bias. This set of particles (277,951) was used for homogeneous refinement with per group global and local CTF refinement to generate the final 3D reconstruction. A custom B-factor of –80 Å^2^ was used for sharpening and the resulting map was used for subsequent model building and refinement steps.

### Single Particle Cryo-EM Structure Model Building and Refinement

AlphaFold3 (50) was used to generate an initial model of the *Tb* H2A.Z-H2B.V NCP with Widom 601 145 bp DNA. UCSF ChimeraX (51) was used to fit the model into the map and to truncate poorly resolved regions (e.g.: histone N-terminal tails). The model was then refined using real space refinement in Phenix (52). A global molecular dynamics simulation was applied to the protein core to reduce artefactual distribution of backbone dihedral angles using ISOLDE (53). Localised simulations were also applied to the C-terminal tail of H2A.Z to improve map to model fit. Iterative adjustments in Coot (54) and ChimeraX were then made to obtain the final model. Validation statistics were obtained using the Phenix implementation of Molprobity (55) and EMRinger (56) and can be found in Supplementary Table 1.

### Micrococcal Nuclease (MNase) Assay

MNase assays were performed essentially as described (19,20). 60 μL reactions containing 1 μg of NCPs and 7.2 units of MNase (New England Biolabs) were incubated at 37°C for 25 min in MNase Buffer (50 mM Tris pH 8.0, 2.5% glycerol (v/v), 25 mM NaCl, 5 mM CaCl2, 1.5 mM DTT). 10 μL were removed from each reaction at 5 min intervals and quenched with 5 μL of Stop Solution (20 mM Tris pH 8.0, 80 mM EDTA, 80 mM EGTA, 0.25% SDS, 0.5 mg.mL^−1^ Proteinase K, New England Biolabs) for 1h at 37°C. Control samples without MNase and the reaction products (44 ng each) were then analysed by native PAGE (5% polyacrylamide, TBE gels stained with Promega Diamond^TM^ Nucleic Acid Dye). Experiments were performed in triplicate.

### Thermal Denaturation Assay

Thermal denaturation assays were performed based on previous studies (19,20,57). NCPs were diluted to 0.5 μM in TDA Buffer (20 mM HEPES pH 7.5, 150 mM NaCl, 1 mM EDTA, 1 mM DTT, 5x SYPRO^TM^ Orange – Invitrogen) in a 50 μL reaction volume and 96-well format. A Biometra TOptical RT-PCR machine was used to heat the plate from 45 to 95°C (0.5°C steps, 30s intervals) and to measure SYPRO^TM^ Orange fluorescence (ex/em = 490/580 nm). Three independent experiments with two technical repeats were performed for each NCP. The technical repeats were averaged and the relative fluorescence intensity (RFU) measured at each temperature point (*x*) was normalised as follows:

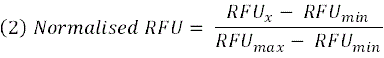

The melting temperature (*T_m_*) values for each NCP were estimated from the peak maximum value of the first derivative of each melting curve using qPCRsoft software (Analytik Jena). The data was visualised using Prism and can be accessed in Supplementary Data File 2.

### DOT1B Methyltransferase Assay

Methyltransferase assays with HMT-DOT1B were performed as described previously(20) using the MTase-Glo^TM^ Kit (Promega) (58). A 2-fold dilution series of 0-500 nM nucleosomes wrapped with Widom 603 193 bp DNA was incubated with 150 nM HMT-DOT1B and 10 μM SAM for 30 min at 37°C in MTase Buffer (20 mM HEPES pH 8.0, 50 mM NaCl, 1 mM EDTA, 3 mM MgCl2, 0.1 mg.mL^−1^ BSA (Thermo Fisher Scientific), and 1 mM DTT) in a 384-well plate format, with two technical repeats for each condition. A standard curve with a 2-fold dilution series of 0-1 μM SAH in MTase Buffer was also prepared. The reactions and standards were then sequentially incubated with 1x MTase-Glo^TM^ Reagent and MTase-Glo^TM^ Detection solution for 30 min each at room temperature. A SpectraMax iD5 plate reader was used to measure the produced luminescence. A linear regression model was fit to the standard curve and the measured relative luminescence units (RLU) from each nucleosome concentration point (*x*) were converted to SAH produced (μM):

Experiments for each nucleosome were performed at least in triplicate. Repeats were averaged, plotted in Prism, and fit to the Michaelis-Menten model. Quantified data is provided in Supplementary Data File 2.

### KKT23 Acetyltransferase Assay

KKT23 acetyltransferase assays were performed based on a previous approach (44), with some modifications. 1 μM NCPs were incubated with 1 μM KKT23 at 37°C in a 50 μL reaction volume in HAT Buffer (50 mM Tris pH 8.0, 50 mM NaCl, 5% glycerol (v/v), 0.1 mM EDTA, 1 mM DTT, 10 mM sodium butyrate, and 0.2 mM acetyl-CoA). After 10, 20, 40, and 60 min, 10 μL were removed from each reaction and quenched by adding 10 μL of 2xSDS loading dye and heating to 95°C for 3 min. The same treatment was applied to control samples without enzyme. In the case of *Tb* canonical NCPs, KKT23 activity was tested at two enzyme concentrations (1 μM and 12.8 μM) and only sampled after 60 min of incubation. After quenching, 8 μL of each sample were loaded on an mPAGE® 4-20% Precast Gel (Millipore).

Western blotting was performed by transferring the proteins onto a methanol-activated PVDF membrane using the Trans-Blot® Turbo^TM^ System mixed MW transfer protocol (Bio-Rad). The PVDF membrane was pre-blocked in 5% milk in PBS-0.01% Tween-20 (PBST) for 30 min at room temperature, washed with PBST three times (5 min each), and incubated with a primary pan-acetyl lysine rabbit antibody (Ac-K^2^-100, 9814, Cell Signaling Technology) overnight at 4°C using a 1:1,000 (v/v) dilution in 5% milk PBST. The membrane was then washed with PBST three times (5 min each) and incubated with a horseradish peroxidase-conjugated anti-rabbit secondary antibody (PI-1000, Vector Labs) for 2 h at room temperature using a 1:5,000 (v/v) dilution in 5% milk PBST. The membrane was washed with PBST three times (5 min each) and incubated with 2 mL of SuperSignal^TM^ West Pico PLUS Chemiluminescent substrate (Thermo Fisher Scientific). Chemiluminescence was detected using a ChemiDoc^TM^ Imaging System (Bio-Rad).

### Magnesium Precipitation of Nucleosome Arrays

Magnesium precipitation assays with non-fluorescent nucleosome arrays were performed as described previously (20). Briefly, 10 μL of 80 ng.μL^-1^ arrays (60 nM) were mixed with 1 volume of 2x Precipitation Buffer (2x MgCl2, 15 mM HEPES pH 7.5, 1 mM EDTA, 1 mM DTT) for 15 min on ice and centrifuged for 15 min at 17,000 x g, 4°C. The concentration of soluble arrays was measured in triplicate based on DNA using a NanoDrop. The data was fit to a non-linear EC50 shift model in Prism.

Magnesium precipitation assays with Alexa647-labeled fluorescent nucleosome arrays were performed as described above, with some modifications. After centrifugation, 12 μL of soluble arrays were diluted 10-fold in Array Storage Buffer (15 mM HEPES pH 7.5, 25 mM NaCl, 1 mM EDTA, 1 mM DTT) and 30 μL of each array (3 nM) were transferred to a 384-well plate in triplicate. Fluorescence was measured on a SpectraMax iD5 instrument (ex/em λ = 630/670 nm).

### Flow Induced Dispersion Analysis

Biophysical analysis of nucleosome-OG488 labelled arrays was performed on a FidaBio Neo instrument using an excitation wavelength of 480nm. 30nM of *Tb* H2AK120C-OGG488 and *Tb* H2A.ZK169C-OGG488 arrays were applied to coated 75 μm capillaries, surrounded by Array Storage Buffer (15 mM HEPES pH 7.5, 25 mM NaCl, 1 mM EDTA, 1 mM DTT). For magnesium titration experiments, Array Storage Buffer was supplemented with a dilution series of magnesium (0-6 mM) and mixed on a capillary within the instrument. The experiment was performed at 25°C and repeated at 200mBar, 140 mBar and 70 mBar, with similar results. The resultant Taylorgrams were fitted using standard settings in FIDA analysis software (version 3.2) showing good polydispersion index, spike counter and signal to noise ratio in the absence of magnesium. Hydrodynamic radius is reported based on repeated measurements in Array Storage Buffer with R^2^ values of fit greater than 0.999. At higher magnesium concentrations, arrays no longer exhibited standard Taylor dispersion even at low system pressures suggesting that the particles were greater than 100nm.

### Fluorescence Microscopy

Fluorescence microscopy was performed with 1/12 Alexa647-labelled nucleosome arrays. 80 ng.μL^-1^ arrays (60 nM) were mixed with 1 volume of 2x Precipitation Buffer (0 or 5 mM MgCl_2_, 15 mM HEPES pH 7.5, 1 mM EDTA, 1 mM DTT) and incubated for 15 min on ice. 10 μL of the arrays were applied to an uncoated microscope slide and imaged on a Zeiss Axio Imager A2 using a 100x/1.4 NA objective and Cy5 filter set. Images were analysed in ImageJ (59) and contrast enhanced equally.

### Affinity Purification (AP) of Nucleosome Interactors

Trypanosome cell extract was obtained from *T. brucei brucei* Lister 427 procyclic cells grown at 37°C to ∼1×10^7^ cell mL^-1^ in SDM-79 medium supplemented with 10% Fetal Calf Serum. Cells were lysed by gentle sonication and dounce homogenisation in Extract Buffer (50 mM Tris pH 8.0, 150 mM NaCl, 10% glycerol (v/v), 2 mM DTT, 0.2% NP-40 (v/v), 1 mM AEBSF,2.2 mM PMSF, 2 mM benzamidine-HCl, 2 μM leupeptin, 1 μg.mL^-1^ pepstatin A). The total protein concentration of the extract (19.5 mg.mL^-1^) was determined using a Pierce^TM^ BCA Protein Assay Kit (Thermo Fisher Scientific) and used as Input in mass spectrometry experiments.

Pulldowns were performed in a similar manner to previous studies (60) and utilised nucleosomes wrapped with biotinylated Widom 601 175 bp that were either pre-treated with glutaraldehyde (+GA) or untreated (-GA). The nucleosomes were crosslinked with 0.0075% (v/v) GA in Nucleosome Storage Buffer (15 mM HEPES pH7.5, 25 mM NaCl, 1 mM DTT) for 5 min on ice and quenched with 45 mM Tris pH 7.5. Non-crosslinked samples were treated in an equivalent manner. 30 μg of each nucleosome sample were then immobilised on ∼30 μL of Streptavidin Sepharose^TM^ High Performance beads (Cytiva) overnight at 4°C in Pulldown Buffer (20 mM HEPES pH 7.9, 150 mM potassium acetate, 10% glycerol (v/v), 1 mM EDTA, 1 mM DTT) supplemented with 0.01% NP-40 (v/v). Subsequent centrifugation steps with the beads were performed for 2 min at 1,500 x g, 4°C.

The beads were washed twice with 500 μL of Pulldown Buffer supplemented with 0.01% NP-40 (v/v) and once with 500 μL of Pulldown Buffer supplemented with 0.01% NP-40 (v/v) and 0.2 mM PMSF (5 min each at 4°C). Following the final wash, the beads were divided into three equal volumes (10 μg nucleosomes per repeat). Each sample was incubated with 500 μL of 3.0 mg.mL^-1^ trypanosome cell extract (1.5 mg protein) for 2h at 4°C. The beads were then washed five times for 10 min each and with 500 μL buffer. The first two washes were performed with Pulldowns Buffer supplemented with 0.01% NP-40 (v/v), the third wash was performed with Pulldown Buffer, and the fourth and fifth washes were performed with Pulldown Buffer prepared without glycerol.

Bound proteins were eluted with 50 μL of Elution Buffer (2M Urea, 100 mM Tris pH 7.5, 10 mM DTT) for 20 min in a ThermoMixer (Eppendorf) at 25°C, 1500 rpm. The samples were alkylated with 6 μL 500 mM iodoacetamide in 100 mM Tris pH 7.5 for 10 min at 25°C, 1500 rpm in the dark. 30 μL of 10 ng. μL^-1^ trypsin (Thermo Fisher Scientific) dissolved in 50 mM acetic acid were then added and incubated for 2h at 25°C, 1500 rpm. The reactions were centrifuged and the supernatants were transferred to fresh tubes. To maximise elution, 50 μL of Elution Buffer were added to the remaining beads for 5 min at 25°C, 1500 rpm. The beads were centrifuged and the supernatants from the first and second elution steps were combined. 15 μL of 10 ng μL-1 trypsin (TFS) in 50 mM acetic acid were added and the reactions were incubated overnight at 25°C, 1500 rpm. 5.5-7.5 μL of 10% TFA (v/v) were added to each sample for sample acidification (pH ≤ 2). 80 μL of each sample were loaded on stage tips prepared with Empore C18 disks as described previously (61).

### Mass Spectrometry (MS) Analysis of Nucleosome Interactors

Pulldown samples were analysed by LC-MS/MS using an Orbitrap Fusion™ Lumos™ Mass Spectrometer (Thermo Fisher Scientific) coupled to an Ultimate 3000 HPLC (Dionex, Thermo Fisher Scientific). A 50 cm (2 μm particle size) EASY-Spray column assembled on an EASYSpray source (Thermo Fisher Scientific) operated at 50°C was used to separate the peptides (Mobile Phase A: 0.1% formic acid, Mobile Phase B: 0.1% formic acid, 80% acetonitrile). The following steps were then performed: peptide loading (0.3μL min-1) and peptide elution (0.25 μL min^-1^) using 2-40% Mobile Phase B for 150 min and 40-95% Mobile Phase B for 11 min. Mobile Phase B was then held at 95% for 5 min and reduced to 2% for the remainder of the run (total run time = 190 min). The MS1 survey scan was performed with a mass resolution of 120,000, scan range of 350-1650 m/z, Automatic Gain Control (AGC) ion target of 5.0×10^6^, and injection time of 20 ms. The MS2 data independent acquisition (DIA) was performed with a mass resolution of 30,000, scan range of 200-2000 m/z, AGC ion target of 3.0×10^6^, and maximum injection time of 55 ms. Higher energy collisional dissociation (HCD) fragmentation (62) was performed with stepped collision energies of 25.5, 27 and 30. We used variable isolation windows throughout the scan range ranging from 10.5 to 50.5 m/z. Narrower isolation windows (10.5-18.5 m/z) were applied from 400-800 m/z and then gradually increased to 50.5 m/z until the end of the scan range. The default charge state was set to 3. The inclusion mass list and corresponding isolation windows are available in Supplementary Data File 3. Data for both MS1 and MS2 were acquired in profile mode.

### AP-MS Data Processing

Raw DIA files were processed using DIA-NN v1.9.2 (63). Identified proteins were searched against the *T. brucei brucei* (strain 927/4 GUTat10.1) reference proteome in UniProt (December 2024 release). A spectral library containing 9,410 proteins was generated automatically using the deep-learning based spectra, retention times (RTs), and ion mobilities (IMs) method. Protease and cleavage settings were specified to trypsin and a maximum of two missed cleavages. Modifications that were specified were carbamidomethylation (fixed), methionine oxidation (variable), and N-terminal acetylation (variable) or lysine acetylation (Supplementary Figure S13). Precursor false discovery rate was set to 1%. Other DIA-NN software parameters were used with default settings. Protein annotations were added using the FASTA database.

A total of 3,575 proteins were identified across all the pulldown samples, of which 3,125 could be confidently detected with ≥ 2 unique peptides and were used for further analysis with DIA-Analyst (Monash Proteomics, https://analyst-suites.org/apps/dia-analyst/). The raw data for each protein can be accessed in Supplementary Data File 3. Next, 3,048 proteins were retained after automatic pre-filtering in DIA-Analyst, which removes proteins with a high number of missing values. Perseus-type imputation was used and P-values were calculated using default settings, which utilise per protein linear model fitting coupled to empirical Bayes statistics from the limma Bioconductor package (64). Functional annotations such as cellular localisation, protein length, ortholog count and others were retrieved from TriTrypDB (35) and used to further filter and prioritise hits. The statistical analyses from DIA-Analyst and annotations for each protein can be accessed in Supplementary Data File 3. The data was visualised using the R package ggplot2.

### Mapping Histone Acetylation in AP-MS Data

The AP-MS dataset was queried for acetylation (UniMod:1) using DIA-NN v1.9.2 and filtered for peptides with a Peptidoform Q-value < 0.01 across all conditions and repeats. Overall, 223 different histone peptides were found, of which 18 were acetylated and revealed 14 modified sites. Percentage acetylation was calculated by dividing the intensity of each peptide by the total intensity of peptides found for a given histone in each repeat.

## Results

### Formation of *T. brucei* H2A.Z-H2B.V nucleosome core particles (NCPs)

H2A.Z is readily identifiable in *T. brucei* (*Tb*) (65), sharing more similarity to H2A.Z orthologs than to canonical H2A histones (Supplementary Figure S1A). However, *Tb* H2A.Z contains several different regions relative to H2A.Z from other species, particularly within the N-terminal tail, L1 loop, and C-terminal tail (Figure 1A, Supplementary Figure S2). *Tb* H2A.Z functions together with the kinetoplastid-specific histone variant H2B.V, which lacks clear homologues outside kinetoplastids (65) and shares only ∼49% sequence similarity to canonical *Tb* H2B. Compared to a set of previously studied H2B variants from mammals (66), plants (67,68), and apicomplexan parasites (69), kinetoplastid H2B.V is among the most different (Supplementary Figure S1B). H2B.V contains a unique C-terminal extension required for viability (70) (Figure 1A), further highlighting its divergence from canonical histones. Even subtle sequence variation within H2A.Z has been shown to substantially alter nucleosome behaviour in other systems (11,33,34), motivating direct biochemical and structural analysis of the trypanosome variants.

**Figure 1:**
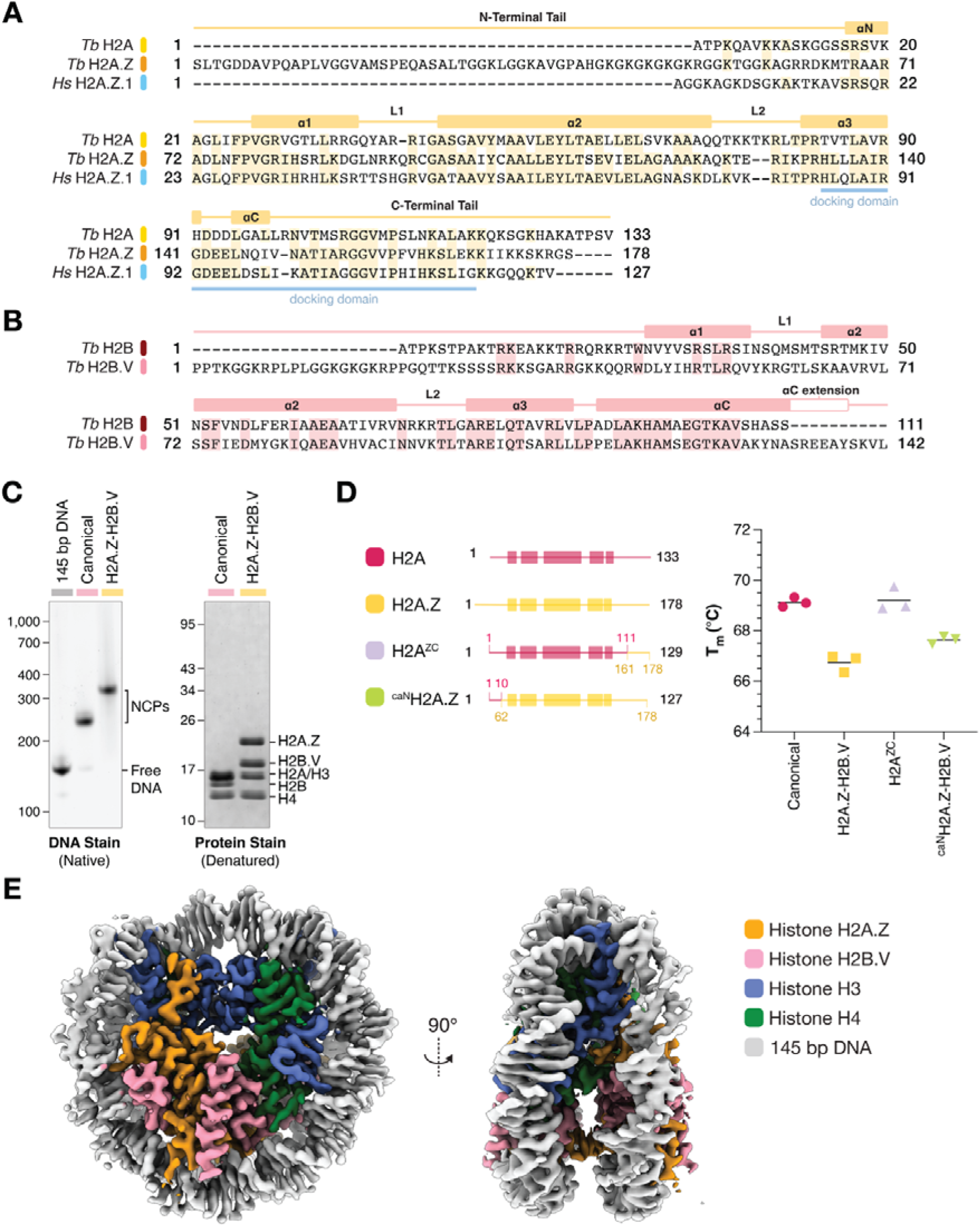
*T. brucei* histone variants H2A.Z and H2B.V form an obligate dimer and assemble into unique nucleosome core particles (NCPs) A. Structure-based multiple sequence alignment of *T. brucei* (*Tb*) H2A, *Tb* H2A.Z, and *H. sapiens*(*Hs*) H2A.Z.1. B. Structure-based pairwise sequence alignment of *Tb* H2B and *Tb* H2B.V. In both A. and B., histone secondary structure is indicated below each alignment and is derived from the structure of the *Tb* H2A.Z-H2B.V NCP. C. Representative native PAGE (left) and SDS PAGE (right) gels of *Tb* Canonical and H2A.Z-H2B.V NCPs reconstituted with Widom 601 145 bp DNA. D. Melting temperature (T_m_) values derived from thermal denaturation profiles of different NCPs (n = 3, see Supplementary Figure S3F–H). The mean T_m_ value for each NCP is indicated with a black line. H2A^ZC^ NCPs were reconstituted with canonical histone H2A fused to the C-terminal tail of H2A.Z (H2A aa 1-111 + H2A.Z aa 161-178). ^caN^H2A.Z NCPs were reconstituted with histone H2A.Z fused to the N-terminal tail of canonical H2A (H2A aa 1-10 + H2A.Z aa 62-178). The NCPs were reconstituted with Widom 601 145 bp DNA. E. Two views of the single particle cryo-EM density map of the H2A.Z-H2B.V NCP coloured based on histones and DNA.

To understand the effect of *Tb* H2A.Z and *Tb* H2B.V on nucleosome structure and function we reconstituted H2A.Z-H2B.V nucleosome core particles (NCPs). We previously purified canonical trypanosome histones from *E. coli* (19) and H2B.V could be similarly purified under standard conditions. However, H2A.Z required an N-terminal 6xHis-TEV tag to improve expression and yield. The tag was removable and had no detectable effect on NCP assembly or properties (Supplementary Figure S3E). Yields for refolding of H2A.Z-H2B.V octamers (with *Tb* H2A.Z, H2B.V, H3, and H4) were low (Supplementary Figure S3A-D), suggesting that complex formation between H2A.Z-H2B.V dimers and the H3-H4 tetramer is unstable. To form wrapped NCPs, we leveraged an approach used previously for unstable H2A.Z nucleosomes (32), mixing H3-H4 tetramers, H2A.Z-H2B.V dimers, and strong nucleosome-positioning Widom 601 145 bp DNA prior to salt dialysis. This yielded homogeneous particles indistinguishable from those prepared with histone octamers (Supplementary Figure S3E). Notably, H2A.Z-H2B.V NCPs displayed reduced electrophoretic mobility than canonical NCPs on native gels, in line with their higher molecular weight (∼15.4 kDa larger) and altered physicochemical properties (Figure 1C).

### *Tb* H2A.Z-H2B.V NCPs are less stable

Consistent with unstable octamer formation, *Tb* H2A.Z-H2B.V NCPs had substantially lower thermal stability than canonical *Tb* NCPs (Figure 1D, Supplementary Figure S3F-H). Their stability was also much lower than both canonical and H2A.Z-containing *H. sapiens* NCPs (33,71) (Supplementary Figure S3G-H)*. Tb* H2A.Z has an unusually long and compositionally bipartite disordered N-terminal tail, comprising an acidic segment followed by a lysine/glycine-rich region that is hyperacetylated *in vivo* (25,72) (Supplementary Figure S2A). Additionally, *Tb* H2A.Z also has a divergent C-terminal tail (Figure 1A), a region that is implicated in nucleosome stability in metazoa (31,73). To test if the N– and C-terminal H2A.Z tails control the stability of H2A.Z-H2B.V NCPs, we performed tail-swapping experiments. Neither the replacement of the H2A.Z N-terminal tail with the canonical tail (^caN^H2A.Z) nor the C-terminal tail swap, replacing the H2A C-terminal tail with the H2A.Z C-terminal tail (H2A^ZC^), were sufficient to account for the reduced stability of H2A.Z-H2B.V NCPs (Figure 1D, Supplementary Figure S3F and S3H). This suggests that nucleosome instability is encoded primarily within the variant histone fold, rather than the unstructured tails.

### Cryo-EM structure of the *Tb* H2A.Z-H2B.V NCP

To gain a detailed understanding of the unique structural features of H2A.Z and H2B.V NCPs, we analysed the complex by single particle cryogenic electron microscopy (cryo-EM). Following mild chemical crosslinking, we obtained a 2.77 Å reconstruction (Figure 1E, Supplementary Figure S4 and S5). This enabled model building of all four core histones and 143 bp of DNA, with only the terminal disordered histone tails unresolved (Supplementary Table S1, Supplementary Figure S5D). The H2A.Z-H2B.V NCP forms a characteristic coin-like shape with central histones wrapping and compacting DNA in a left-handed supercoil, reminiscent of structures from both *T. brucei* and other species NCPs (19,30,74) (Figure 1E).

Overall, the structure reveals a combination of unique and conserved features at the secondary structure and amino acid level (Figure 2A). For example, the C-terminal extension of H2B.V (aa 133-142) leads to an ordered extension of the αC helix (aa 133-138) (Figure 2B) uncommonly observed in NCP structures (75). The extension covers the DNA gyre near super-helical location (SHL) 4.0 and contains an extra tyrosine (Y138) and arginine (R134) that likely reinforce DNA binding. At the dimer–tetramer interface, a hydrogen bonding network that comprises H2B-K74, H2B-E81, and H4-Y73 in *T. brucei* canonical histones is ablated with a substitution of a hydrophobic valine residue (H2B-K74 to H2B.V-V95; Supplementary Figure S6A). This point of difference is conserved across diverse kinetoplastids (Supplementary Figure S2B), indicating evolutionary remodelling of octamer connectivity.

**Figure 2:**
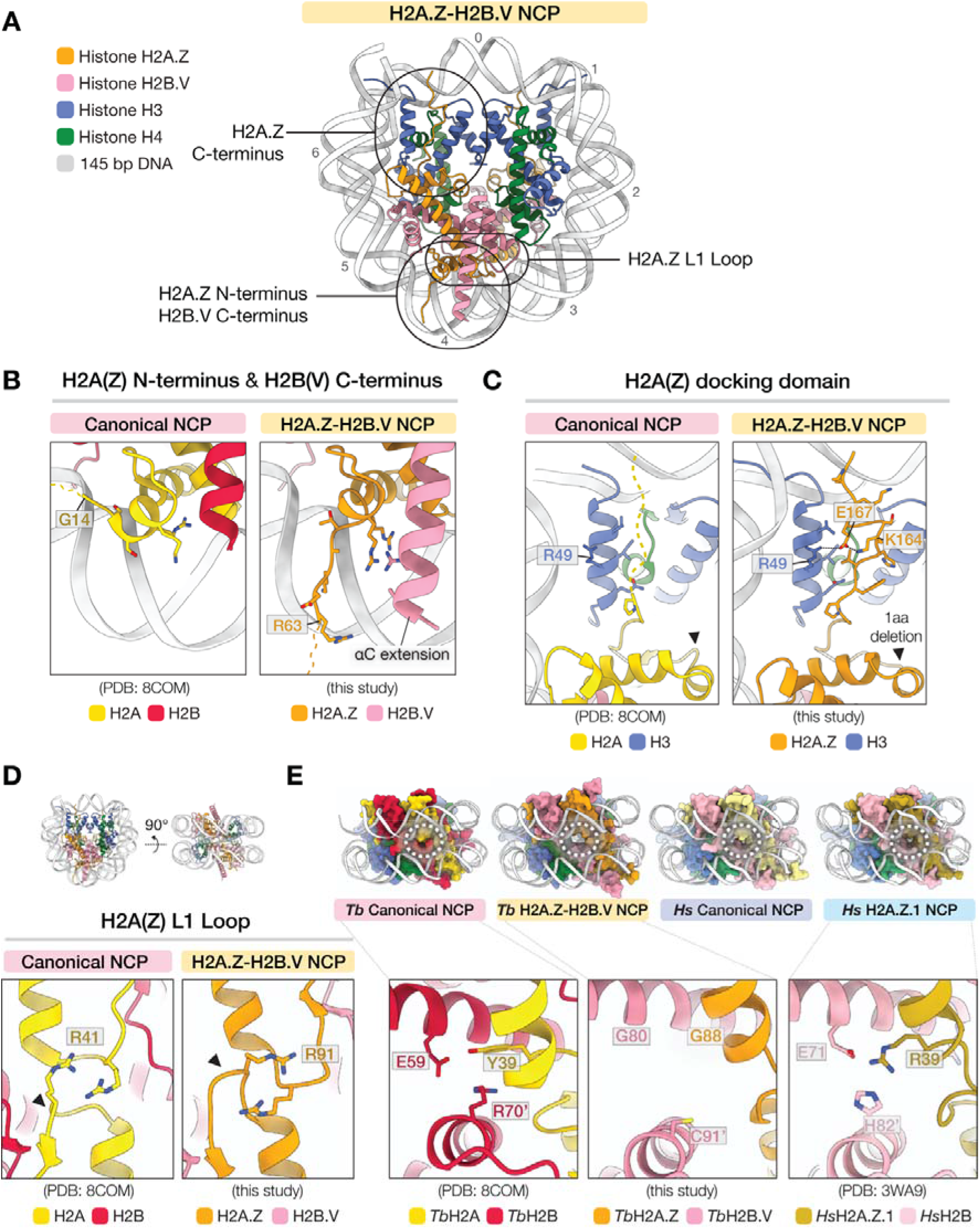
The structure of the *T. brucei* H2A.Z-H2B.V NCP reveals both conserved hallmarks of eukaryotic H2A.Z as well as trypanosome-specific features. A. Overview of the *T. brucei* H2A.Z-H2B.V NCP model highlighting regions of H2A.Z and H2B.V that are magnified regions in panels B-D. Superhelical locations (SHL) on the DNA are shown. B. Magnified view of the H2A N-terminal tail and the αC helix of H2B (left PDB: 8COM (19)) compared to the equivalent region in H2A.Z-H2B.V (right). Better ordering of the H2A.Z N-terminal tail and a structured extension of the H2B.V αC helix lead to increased DNA-histone interactions. C. Magnified view of the *Tb* canonical (left, PDB: 8COM (19)) and H2A.Z (right, this study) docking domain region between histones H2A(Z) and H3, with selected residues shown and highlighted. Better ordering of this region in H2A.Z allowed us to build into the density. D. Enlarged view of the H2A(Z) L1 loop dimerization interface comparing differences in loop structure in this region. E. Top: Comparison of different NCP structures (*T. brucei* canonical, PDB: 8COM (19); *T. brucei*H2A.Z-H2B.V, this study; *H. sapiens* canonical, PDB: 7XD1 (80); *H. sapiens* H2A.Z.1, PDB: 3WA9 (34)) showing distinctive gaping and a void at SHL 5 highlighted with a white dashed oval. Histones are shown in surface representation and DNA is shown as a cartoon. Bottom: Magnified views of the H2A.Z void region.

The structural hallmarks of eukaryotic H2A.Z are conserved. The *Tb* H2A.Z docking domain (Figure 1A, blue line) adopts a conserved fold that packs against the αN helix of histone H3 (Figure 2C). The base of the docking domain includes a conserved, single amino acid deletion between H2A.Z-V149 and N150 that was shown to control chaperone specificity for H2A.Z over H2A in other eukaryotes (76) (Figure 2C). In contrast, the interaction network between *Tb* H2A.Z and H3 is slightly altered, including the replacement of a conserved hydrophobic residue (I118 in human H2A.Z) (Figure 1A) with a glutamate (E167 in *Tb* H2A.Z). The glutamate likely forms interactions with both *Tb* H3-R49 and *Tb* H2A.Z-K164 (Figure 2D). These residues are broadly conserved across kinetoplastids (Supplementary Figure S2A) and likely stabilise the docking domain and associated DNA, explaining the better ordering of this region in our cryo-EM density compared to canonical *Tb* H2A (Figure 2C).

As in other eukaryotic H2A.Z NCP structures (11,30,31), the *Tb* H2A.Z L1 loop is longer and forms a different L1-L1’ interaction interface compared to canonical H2A (Figure 2D). Notably, this interface is larger than in previously characterised H2A.Z nucleosomes and is primarily stabilised by reciprocal packing of Arg-91 side chains across the two *Tb* H2A.Z protomers. Equivalent arginine residues are also present in the *Tb* Canonical NCP but are missing in H2A L1 loops from other eukaryotes, where they are replaced by serines. Substituting the serines for arginines at this position was previously shown to stabilise H2A.Z-DNA interactions (33). In the *Tb* H2A.Z-H2B.V NCP, the Arg-91 side chains do not appear to bind to the nearby DNA, but map interpretation is partially limited by lower resolution in this region.

Interestingly, the nucleosome DNA exhibits gaping near the L1-L1’ interface, with larger separation between the two gyres of DNA (Figure 2E). Inter-gyre gaping phenomena have previously been suggested to be more pronounced for other H2A.Z nucleosomes (77–79), but appear amplified in the *Tb* H2A.Z-H2B.V NCP structure (Supplementary Figure S6B). The opening correlates with expansion of a solvent-exposed cavity behind the L1-L1’ loop, which normally contains conserved, bulky residues, present in both *Tb* H2A and human H2A.Z (*Tb* H2A-Y39, *Hs* H2A.Z-R39). Instead, *Tb* H2A.Z contains a smaller glycine residue (G88). H2B.V-specific alterations for smaller residues also contribute to this void (*Tb* H2B.V-G80 vs. *Tb* H2B-E59/*Hs* H2B-E71 and *Tb* H2B.V-C91 vs. *Tb* H2B-R70/*Hs* H2B-H82) (Figure 2E). Additionally, the H2B.V N-terminal tail, which is proximal to the cavity, is less ordered as it exits the nucleosome. Overall, the amino acid changes that contribute to this cavity are conserved in *Trypanosoma* species (Supplementary Figure S2) and may help explain the lower stability of H2A.Z-H2B.V nucleosomes.

### H2A.Z-H2B.V form obligate dimers in nucleosomes due to steric constraints

H2A.Z and H2B.V have been mapped to the same genomic locations in *T. brucei* (14,23) and co-immunoprecipitate with each other but not canonical histones (23,25), suggesting they form obligate dimers. Indeed, when we attempted to assemble heterotypic H2A.Z-H2B and H2A-H2B.V mixed dimers both canonical-variant combinations were susceptible to high levels of aggregation compared to canonical or variant combinations alone (Supplementary Figure S6C-D). Structural analysis of the *Tb* H2A.Z-H2B.V interface reveals multiple steric incompatibilities that would prevent productive pairing with canonical H2A or H2B (Supplementary Figure S6E-F). The residues responsible for this incompatibility are broadly conserved across trypanosomatids (Supplementary Figure S2), indicating co-evolution of the histone pair to ensure dimerisation.

### H2A.Z-H2B.V NCPs have altered DNA binding properties that contribute to their instability

In addition to reduced protein-protein interfaces, *Tb* H2A.Z-H2B.V NCPs are more prone to salt-induced disassembly (Figure 3A), suggesting weaker electrostatic interactions with the DNA phosphate backbone. DNA-protein interactions are altered at multiple positions around the nucleosome core, but most are locally compensatory, leading to a similar number of side chain-DNA backbone interactions compared to the canonical *Tb* nucleosome. However, the *Tb* H2A.Z-H2B.V NCP features several differences at SHL 3.5 (Figure 3B). Compared to the canonical H2A-H2B dimer, H2A.Z and H2B.V have a reduced number of positively charged residues in this region, due to three arginine-to-lysine substitutions, and a negatively charged residue (H2A.Z-D87) (Figure 3B). We previously showed that the cluster of positive charges at SHL 3.5 in *Tb* canonical NCPs has functional consequences, serving as a critical roadblock for Micrococcal Nuclease (MNase) digestion (19). Indeed, MNase digestion of canonical NCPs leads to a ‘stall’ band that we previously mapped to the SHL 3.5 region (Figure 3C). In contrast, H2A.Z-H2B.V NCPs are digested faster and have a smear-like MNase digestion pattern with higher accumulation of further digestion products (Figure 3C), suggesting a more dynamic range of digestion outcomes, perhaps derived from both sides of the NCP. These altered patterns in DNA binding may have functional consequences on how proteins such as RNA polymerases access nucleosome-bound DNA in H2A.Z-H2B.V-enriched chromatin.

**Figure 3:**
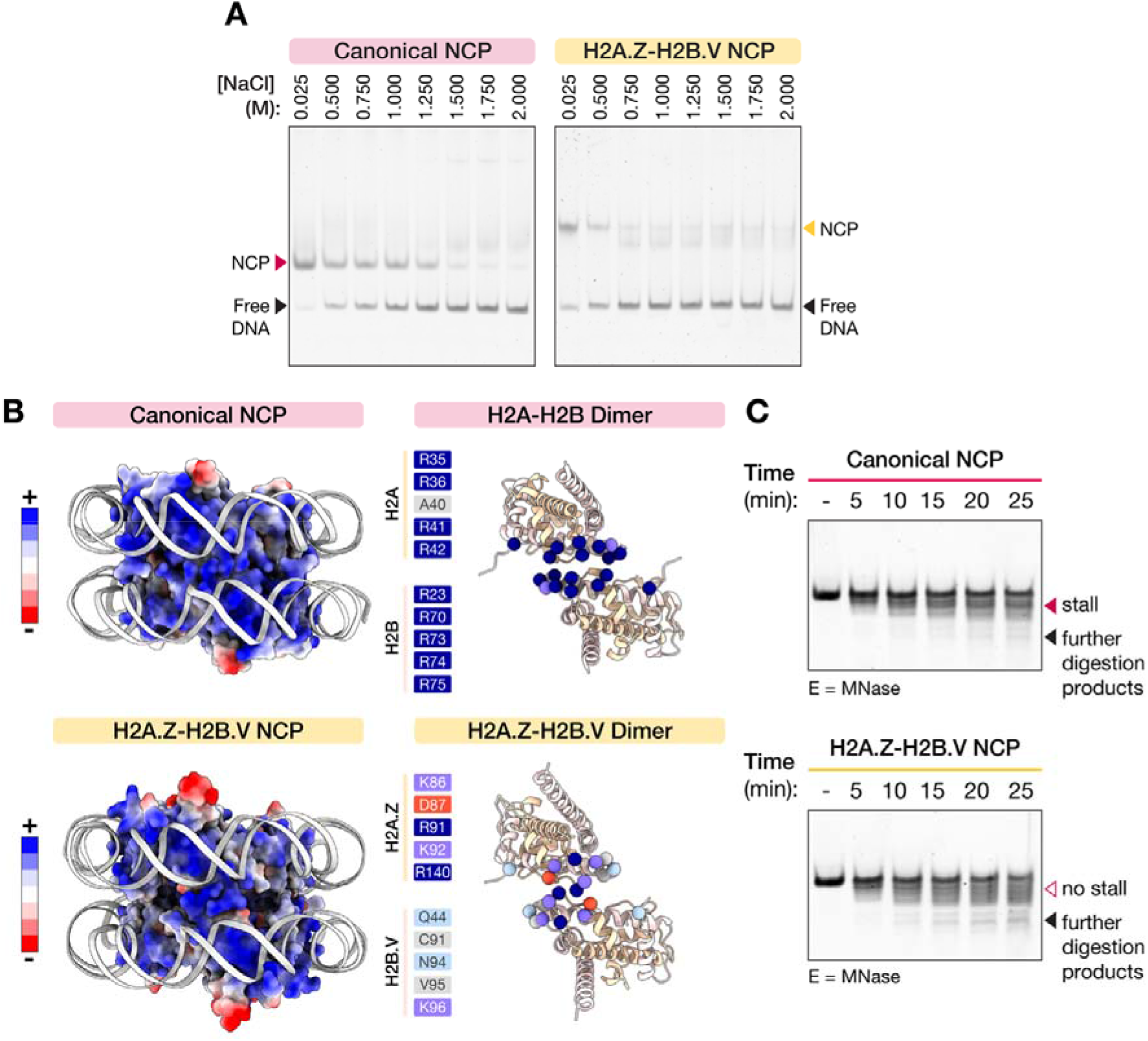
The histone core in H2A.Z-H2B.V NCPs has altered DNA binding properties compared to canonical NCPs. A. Representative gels from salt stability assays, where canonical and H2A.Z-H2B.V NCPs were treated with increasing concentrations of NaCl prior to separation by native gel electrophoresis (n = 3). B. Left: The structure of the canonical NCP (PDB: 8COM (19)) and H2A.Z-H2B.V NCP (this study) shown at super-helical location (SHL) 3.5. Histone octamers are coloured by surface electrostatic potential and DNA is shown as a grey cartoon. Right: Cartoon representation of the H2A-H2B dimer (PDB: 8COM (19)) and H2A.Z-H2B.V dimer (this study) at SHL 3.5. Residues that contribute to the SHL 3.5 interface are shown as spheres and listed on the left in corresponding colours. C. Representative gels showing micrococcal nuclease (MNase) digestion of canonical and H2A.Z-H2B.V NCPs over time (n = 3). Digestion products that preferentially appear in canonical or H2A.Z-H2B.V samples are indicated (“stall” and “further digestion products”). Lane labelled “-“ corresponds to samples incubated for full timecourse in the absence of MNase enzyme.

### The H2A.Z C-terminal tail is structurally distinct from H2A and acts as an altered enzyme substrate

Given the reduced stability of H2A.Z-H2B.V NCPs (Figure 1D, Figure 3), we were surprised by the good ordering of the H2A.Z C-terminal docking domain, H3 αN helix, and entry/exit DNA relative to the canonical NCP (Figure 2C). 3D classification of the cryo-EM data revealed different amounts of density for this region, where ∼81% of total averaged particles contain an ordered H2A.Z docking domain (Figure 4A). The *Tb* H2A.Z-H2B.V NCP also undergoes a degree of dynamic DNA breathing (∼44% of particles, classes 3-4), but this phenomenon does not seem to be as frequent as observed for the canonical NCP (∼70% of particles (19)), nor as dominant as observed for *Tb* H3.V-H4.V variant NCPs (∼100% of particles, with different levels of unwrapping/splaying (20)) under similar conditions. In other eukaryotes, divergent sequences mapping to the C-terminal tail in H2A or H2A.Z variants have been proposed to modulate DNA entry/exit flexibility (11,31,32,77,81), highlighting how the structural role of this region is repeatedly adapted across different evolutionary contexts.

**Figure 4:**
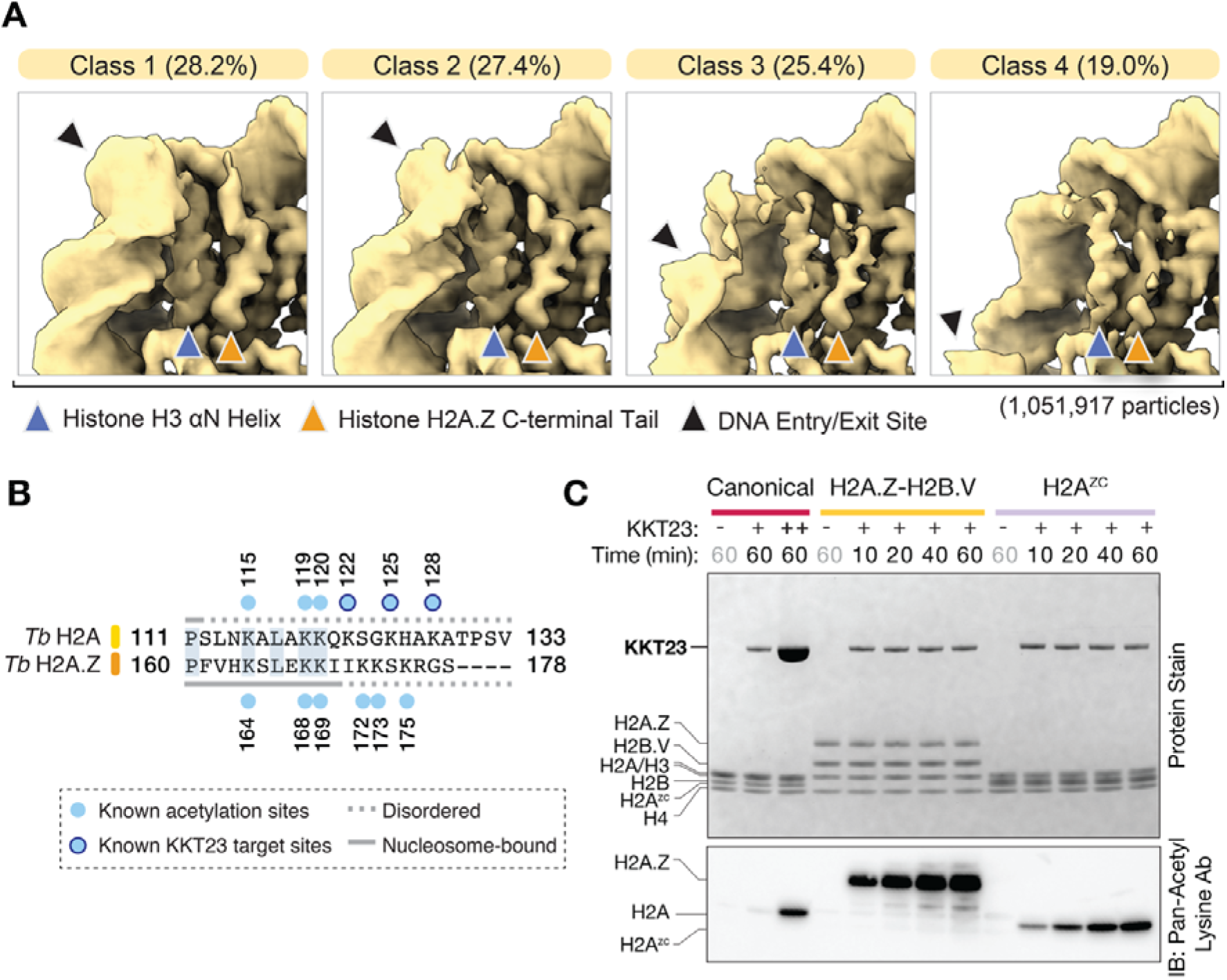
The H2A.Z C-terminal region is ordered in the nucleosome and is acetylated by KKT23. A. 3D classes from cryo-EM data of the H2A.Z-H2B.V NCP. The percentage of particles belonging to each class is provided in each heading (1,055,641 particles in total). The positions of the H3 αN helix, H2A.Z C-terminal tail, and DNA entry/exit site are indicated. B. Pairwise sequence alignment of the C-terminal tail of *Tb* histones H2A and H2A.Z. Known histone acetylation sites from previous studies are highlighted (25,44,72). C. *In vitro* acetylation of NCPs reconstituted with Widom 601 145 bp DNA by the *Tb* histone acetyltransferase KKT23 over time. Left: SDS PAGE gel of the acetylation reactions, right: Western blot probing acetylation. The “-” lanes represent reactions in the absence of KKT23. Canonical NCPs (1 μM) were incubated with 1 μM and 12.8 μM KKT23 and analysed after 60 min (Lanes 2 and 3 respectively). H2A.Z-H2B.V and H2A^ZC^ NCPs (1 μM) were incubated with 1 μM KKT23 and analysed after 10, 20, 40, and 60 min.

Interestingly, the C-terminal tail of both H2A and H2A.Z is highly acetylated in *T. brucei* (25,72) (Figure 4B), adding another layer of functional regulation to this region. Specific sites on the *Tb* H2A C-terminal tail were previously shown to be acetylated by the *Tb* KKT23 acetyltransferase (Figure 4B) (44). Given the differences in the H2A.Z C-terminal tail sequence and structure, we explored whether H2A.Z could be acetylated by KKT23. Surprisingly, after incubation with purified KKT23 and its cofactor acetyl-CoA, robust acetylation signal could be rapidly observed on H2A.Z, surpassing KKT23 activity on canonical H2A (Figure 4C). A tail swap mutant (H2A^ZC^), where the tail region of canonical H2A is swapped for residues from H2A.Z, also led to high acetylation activity indicating that the H2A.Z C-terminal tail is sufficient to confer enhanced KKT23 substrate preference (Figure 4C). H2A.Z C-terminal tail acetylation was reduced upon RNAi knockdown of KKT23 (44), despite H2A.Z-containing nucleosomes not being actively enriched, supporting the H2A.Z C-terminal tail as an additional target for KKT23-mediated acetylation in *T. brucei* parasites.

### H2A.Z-H2B.V incorporation leads to altered properties in nucleosome arrays

Changes to the path of DNA as it exits the nucleosome have functional implications on higher order chromatin folding (20,31,82). H2A.Z and H2B.V are enriched at ∼10 kb transcription start site regions (TSS) (14,15), suggesting that they could form adjacent tandem nucleosome arrays in the genome. To test whether these variants have a direct role on chromatin structure we reconstituted 12-mer arrays with H2A.Z-H2B.V and compared them to canonical arrays (Supplementary Figure S7). Flow induced dispersion analysis (FIDA) of nucleosome arrays revealed that they were typically monodisperse. Notably, the H2A.Z-H2B.V arrays had a larger average hydrodynamic radius (28.1 nm) compared to canonical arrays (21.3 nm) (Figure 5A, Supplementary Figure S8A-B), in line with their proposed open configuration at TSS.

**Figure 5:**
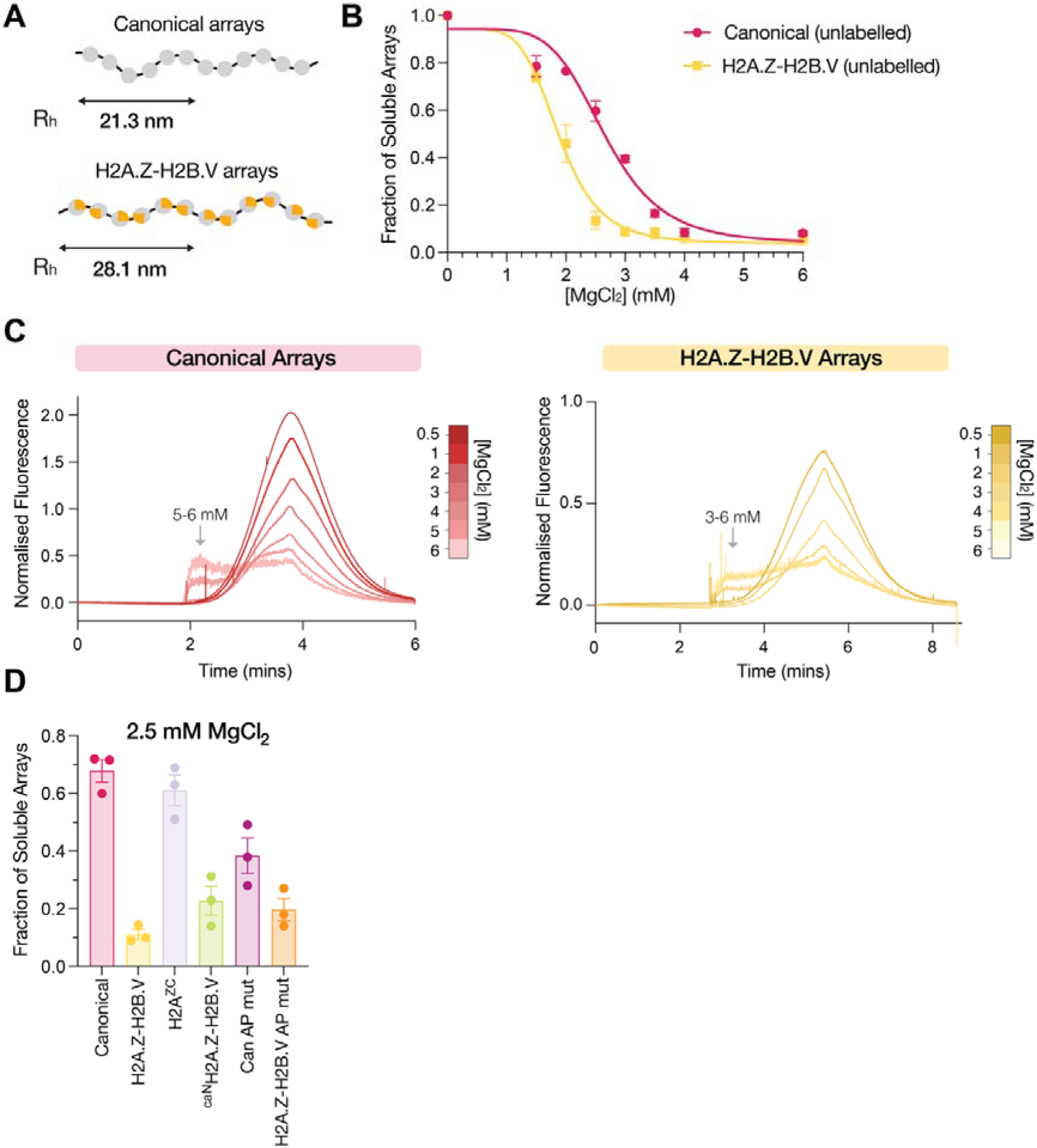
H2A.Z-H2B.V nucleosome arrays are open and prone to self-association. A. The hydrodynamic radius (Rh) of fluorescently labelled nucleosome arrays reconstituted with 12×177 bp Widom 601 DNA calculated from the fit of Flow induced dispersion analysis (FIDA) taylorgram data (n=3). Canonical arrays were labelled with *Tb* H2AK120C-OGG488 and H2A.Z-H2B.V arrays were labelled with *Tb* H2A.ZK169C-OGG488. B. Magnesium precipitation assay with unlabelled canonical and H2A.Z-H2B.V nucleosome arrays reconstituted with 12×177 bp Widom 601 DNA measured by DNA absorbance at A260 (n = 2). A nonlinear EC50 shift model was fit to the data. Error bars represent the standard error of mean (SEM). C. Normalised FIDA Bio Taylorgrams plotting the retention time in the capillary against baseline subtracted fluorescence intensity at 480 nm. An overlay of a MgCl_2_ titration series (0.5-6.0 mM) is shown for both *Tb* H2AK120C-OGG488 arrays (canonical) and *Tb* H2A.ZK169C-OGG488 arrays (H2A.Z-H2B.V). The MgCl_2_ concentrations at which arrays transition to non-Taylor dispersion behaviour and likely form larger particles are indicated with an arrow. Samples were run at different pressures due to different sizes leading to altered retention times of the peaks. D. Magnesium precipitation assay performed using 2.5 mM MgCl_2_ on fluorescently-labelled nucleosome arrays reconstituted with 12×177 bp Widom 601 DNA, 1/12 Canonical-Alexa647 octamers, and 11/12 unlabelled octamers of interest (n = 3). Error bars represent SEM. H2A/H2A.Z tail swaps are defined in Figure 1D. Canonical AP mut = H2A E61A, D93A, D94A H2B D93A; H2A.Z-H2B.V AP mut = H2A.Z E113A, E143A, E144A H2B.V E114A.

The nucleosome arrays were also differentially affected by magnesium cations, which mask DNA backbone charges and promote array self-association. H2A.Z-H2B.V arrays were more susceptible to magnesium precipitation, indicating a higher propensity for self-association (Figure 5B). At higher magnesium concentrations, both canonical and H2A.Z-H2B.V arrays formed particles that were consistent with radii greater than 100 nm in a dose-dependent manner, suggesting inter-array association rather than intra-array condensation (5) (Figure 5C). Again, H2A.Z-H2B.V arrays required lower magnesium concentrations to form array-array interactions. Indeed, fluorescence microscopy analysis revealed the formation of discrete non-spherical aggregates (83) for H2A.Z-H2B.V arrays in the presence of 2.5 mM MgCl_2_, not observed for the canonical arrays (Supplementary Figure S8C).

Based on using chimeric histone nucleosome arrays, increased H2A.Z-H2B.V self-association could not be fully explained by the H2A.Z N– and C-terminal tails (Figure 5D, Supplementary Figure S8C). The nucleosome acidic patch is a solvent-exposed region of negative charge formed between H2A and H2B. In other eukaryotes, the acidic patch has been shown to promote nucleosome-nucleosome interactions in both canonical and H2A.Z arrays (31,84,85). While charge neutralisation of four acidic patch residues resulted in a slight increase in precipitation of canonical arrays, H2A.Z-H2B.V arrays were largely unaffected (Figure 5D, Supplementary Figure S8C). Overall, this suggests that H2A.Z-H2B.V arrays are more open and prone to forming interactions with neighbouring arrays, likely through a combination of nucleosome features. Interestingly, TSS enriched with H2A.Z-H2B.V cluster into distinct domains in the trypanosome nucleus (27). The biophysical features of H2A.Z-H2B.V nucleosomes observed here may directly contribute to this clustering.

### The H2A.Z-H2B.V acidic patch forms a biochemically distinct binding interface

In other eukaryotes, the H2A.Z acidic patch is extended by one residue and this subtle change is sufficient to mediate altered chromatin compaction and chromatin-protein interactions (31,84,86). Although trypanosomatid H2A.Z lacks the additional acidic residue, the electrostatic surface potential of the H2A.Z-H2B.V NCP structure reveals an extended and more open acidic patch relative to canonical *Tb* NCPs (Figure 6A-B). The widening of the patch is likely due to charge neutralisation of H2A.Z-A120 (H2A-K68) and the lengthening of the patch occurs via H2A.Z-D73 substitution (H2A-G22) and the H2B.V αC helix extension (H2B.V-E135, H2B.V-E136, Figure 6C). While core acidic patch residues are conserved across kinetoplastids, these additional features are largely trypanosomatid-specific (Supplementary Figure S2), indicating lineage-restricted changes to this interaction surface.

**Figure 6:**
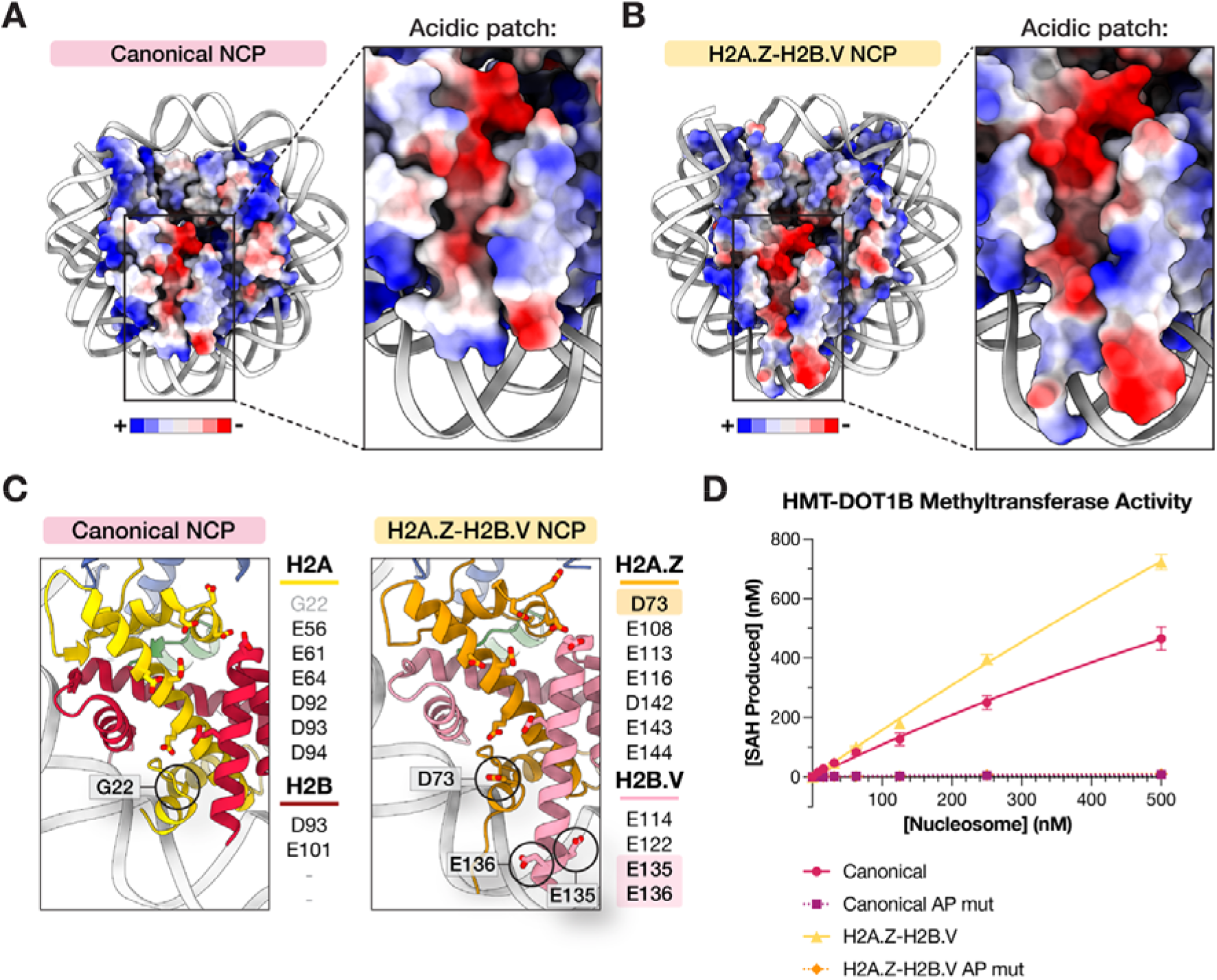
H2A.Z-H2B.V NCPs have a distinct acidic patch that promotes nucleosome interactions. A-B. Structural overview of the canonical acidic patch (panel C, PDB: 8COM) (19) and the H2A.Z-H2B.V acidic patch (panel D, this study). Histone octamers are coloured by surface electrostatic potential and DNA is shown as a grey cartoon. C. A magnified view of residues that form the acidic patch in the canonical NCPs (PDB: 8COM) (19) and H2A.Z-H2B.V NCPs (this study). Non-canonical acidic patch residues from the H2A.Z-H2B.V NCP and residues in equivalent positions in the canonical NCP are circled. D. Methyltransferase assay (MTase-Glo^TM^) with 150 nM His-MBP-TEV (HMT) tagged *Tb* DOT1B. 10 μM co-factor SAM, and increasing concentrations of nucleosomes reconstituted with Widom 603 193 bp DNA (n = 3). The y-axis shows the concentration of the methylation byproduct SAH. The Michaelis-Menten model has been fit to the data. Error bars represent SEM. Canonical AP mut = H2A E61A, D93A, D94A H2B D93A; H2A.Z-H2B.V AP mut = H2A.Z E113A, E143A, E144A H2B.V E114A

To test the functional consequences of this altered surface, we examined the activity of the trypanosome histone methyltransferase DOT1B, which was proposed to engage the nucleosome acidic patch (19,87). We previously purified DOT1B and saw catalytic activity on nucleosomes (20). We now observed that neutralisation of acidic patch charge in the canonical acidic patch (AP mutant) abolishes DOT1B activity *in vitro* (Figure 6D), consistent with acidic patch dependence. Notably, DOT1B displayed a modest increase in activity on H2A.Z-H2B.V nucleosomes, indicating that the altered acidic patch can enhance, rather than disrupt, at least one chromatin-associated enzymatic interaction. Structurally equivalent acidic patch mutations in H2A.Z-H2B.V also abolished activity (Figure 6D). Together, these data define the H2A.Z-H2B.V acidic patch as a biochemically distinct interface that can modulate chromatin enzyme activity.

### H2A.Z-H2B.V nucleosomes have distinct interacting partners

In *T. brucei,* many chromatin proteins have been suggested to localise to H2A.Z-H2B.V-enriched transcription start regions (88–90). Using our purified nucleosomes as baits, we performed an affinity purification mass spectrometry experiment to identify direct components of the H2A.Z-H2B.V interactome (Figure 7A). Given the low stability of the variant nucleosomes, we prepared both canonical and H2A.Z-H2B.V nucleosomes with and without mild chemical crosslinking. Immobilised nucleosomes were incubated with *T. brucei* procyclic cell lysate in triplicate and analysed by DIA LC-MS/MS (Figure 7A, Supplementary Figure S9). Overall, 3,048 unique proteins could be confidently identified across all conditions, enabling robust pairwise comparisons (Supplementary Figure S9B-C).

**Figure 7:**
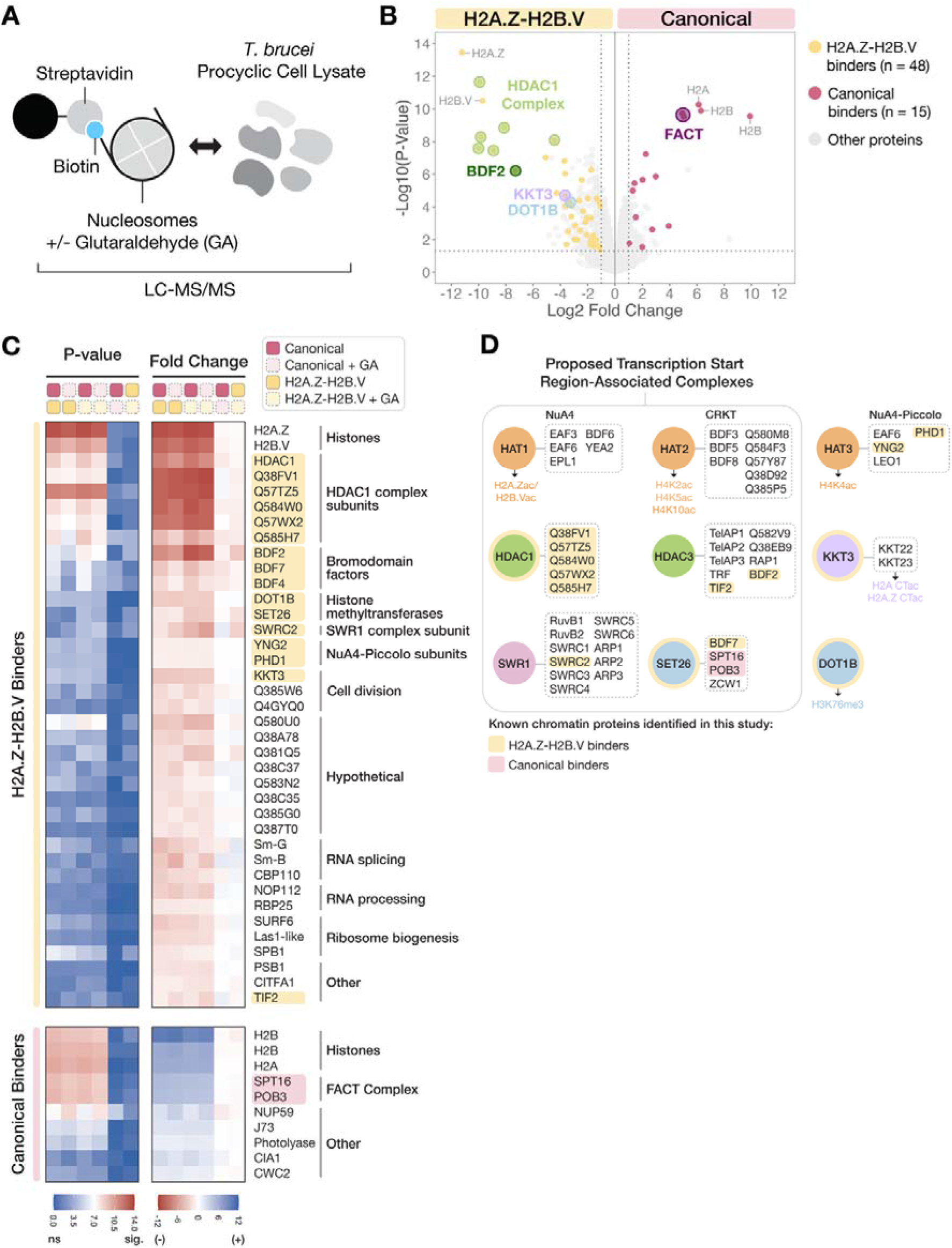
H2A.Z-H2B.V NCPs serve as a direct interaction platform for chromatin proteins at transcription start regions. A. A schematic diagram of the affinity purification-mass spectrometry (AP-MS) approach used to identify nucleosome interactors. Nucleosome reconstituted with biotinylated Widom 601 175 bp DNA were immobilised on streptavidin beads and used to pull down interactors from *T. brucei* Lister 427 procyclic cell lysate. Both native and glutaraldehyde (GA)-stabilised nucleosomes were used (n=3 each). Nucleosome interactors were identified by liquid chromatography tandem mass spectrometry (LC-MS/MS). B. A volcano plot comparing proteins that were preferentially identified in canonical and H2A.Z-H2B.V pulldowns. Proteins that consistently bound to both canonical and canonical+GA nucleosomes are shown in red (15 proteins) and proteins that consistently bound to both H2A.Z-H2B.V and H2A.Z-H2B.V+GA nucleosomes are shown in yellow (48 proteins). Relevant histones and key interactors of canonical and H2A.Z-H2B.V nucleosomes are highlighted if they appear above the P-value threshold of 0.05 (horizontal dashed line) and Log_2_ fold change threshold of |1| (vertical dashed lines). C. Heatmaps showing –log_10_ P-values (left) and Log_2_ fold change values (right) from pairwise comparisons of different nucleosome pulldowns (canonical, canonical+GA, H2A.Z-H2B.V, and H2A.Z-H2B.V+GA). The name or UniProt identifier of each protein is shown on the right. Only proteins with predicted or known nuclear localisation are shown (38/48 H2A.Z-H2B.V nucleosome binders and 10/15 canonical nucleosome binders). In the p-values heatmap, blue tiles represent less significant hits (“ns”) and red tiles represent more significant hits (“sig”). In the fold change heatmap, negative (red) and positive (blue) values were assigned to each condition and are indicated in the heading. D. A summary of how nucleosome interactors identified panel C. overlap with trypanosome chromatin proteins and protein complexes previously shown or predicted to interact with histones (25,26,28,44,95,102–105). H2A.Z-H2B.V nucleosome interactors are highlighted or circled in yellow. Canonical interactors are highlighted in pink. HAT = histone acetyltransferase, HDAC = histone deacetylase, KKT = kinetoplastid kinetochore protein, SWR = SWI2/SNF2-Related (Complex), SET = Su(var)3-9, Enhancer-of-zeste and Trithorax, DOT = disruptor of telomeric silencing. For a full list of protein identifiers from panels B-D, see Supplementary Data File 3.

As expected, H2A.Z and H2B.V were the most enriched proteins in variant pull-downs, whereas canonical H2A and H2B were enriched in control nucleosomes (Figure 7B). Pairwise comparisons identified 48 H2A.Z-H2B.V specific interactors, of which 79% were previously shown to have nuclear localisation (91,92) and included multiple chromatin-associated factors of interest (Figure 7C-D, Supplementary Figure S10-S12). These overlapped with previously defined TSS-associated proteins, particularly Class II factors that colocalise with H2A.Z domains such as BDF2 and SET26 (28) (Figure 7C, Supplementary Figure S11). The enriched factors also included core chromatin regulators such as DOT1B and KKT3, an interactor of KKT23 (93) (Figure 7C, Supplementary Figure S10-S11), suggesting a potential link to our enzymatic assays (Figure 4C, Figure 6D). While we used recombinant unmodified nucleosome baits, many interactors were also enriched for roles in histone acetylation (Figure 7C-D). Acetylation of bait histones was detected during incubation (Supplementary Figure S13), but the acetylation levels were likely too low to explain these enrichment patterns, suggesting direct recognition of the nucleosomal surface rather than acetylation-dependent binding.

SWRC2, a trypanosome homolog of the H2A.Z chaperone YL1 and a subunit of the SWR1 remodelling machinery responsible for H2A.Z deposition (26,28), was also strongly enriched (Figure 7D, Supplementary Figure S11), consistent with direct recognition of the variant nucleosome (76,94). In contrast, both subunits of the FACT histone chaperone complex (95) were depleted from H2A.Z-H2B.V nucleosomes but enriched on canonical particles (Figure 7C, Supplementary Figure S10-S11), suggesting that this chaperone cannot interact with *Tb* H2A.Z-H2B.V. In well studied eukaryotes, FACT has been shown to bind only to H2A-H2B and not H2A.Z-H2B dimers, helping restrict the localisation of H2A.Z on a genome-wide scale (96).

Strikingly, all six subunits of a HDAC1 complex were among the most strongly enriched interactors (28) (Figure 7B-C, Supplementary Figure S10, Supplementary Figure S12A). HDAC1 is essential for parasite viability in both *T. brucei* (97) and *T. cruzi* (98) and has previously been investigated as a potential drug target (99,100). Atypically for a deacetylase, knockdown of HDAC1 was found to promote silencing in *T. brucei*, suggesting an activating role in gene expression (101). Predicted structural similarity analysis of the most significantly enriched subunit (Q57TZ5) revealed potential similarity to SIN3 family scaffold proteins (41), suggesting that the HDAC1 complex could be functionally related to SIN3-HDAC co-repressor/activator complexes (Supplementary Figure S12B-D). Collectively, these data define H2A.Z-H2B.V nucleosomes as a specialised interaction platform for TSS-associated chromatin regulators while excluding some canonical histone-binding machinery.

## Discussion

Our structural and biochemical analysis reveals that *T. brucei* H2A.Z-H2B.V nucleosomes form highly specialised chromatin. Histone-histone and histone-DNA interfaces in H2A.Z-H2B.V nucleosomes are extensively altered compared to both canonical trypanosome histones and H2A.Z from other model organisms (Figures 1-4). These alterations generate nucleosomes that are intrinsically labile (Figure 1 and Figure 3), more open when assembled into nucleosome arrays (Figure 5), and amenable to distinct interactions with chromatin factors (Figure 4 and Figures 6-7). Overall, these findings provide a mechanistic basis for variant definition of transcription start regions (TSSs) in trypanosomatids.

H2A.Z and H2B.V form an obligate heterodimer and are incompatible with canonical histone pairing *in vitro* (Supplementary Figure S6C-F), providing a structural explanation for the co-localisation and interdependence observed *in vivo* (14,15,23). Similar obligate pairing of H2A.Z with a specialised H2B variant has also evolved in apicomplexan parasites (69,106) and may represent a general mechanism to prevent combinatorial dilution of histone properties, instead enforcing a robust variant state that is deposited and maintained as a single functional unit. Consistent with this, FACT and SWRC2 show mutually exclusive preferences for canonical and H2A.Z-H2B.V nucleosomes in our proteomic screen (Figure 7). The kinetoplastid-specific H2B.V C-terminal αC extension is essential for parasite viability (70). We find it simultaneously adds DNA contacts and extends the acidic patch region, coupling DNA wrapping and effector recruitment in one short structural alteration.

H2A.Z-H2B.V nucleosome instability in *T. brucei* is amplified beyond that reported for other H2A.Z nucleosomes (8,34,73), reflecting a combination of weakened histone-histone contacts, reduced histone-DNA interactions and an enlarged internal cavity (Figures 1-3). Instability and openness have been proposed as an inherent mechanism for genome accessibility (107,108). Accordingly, H2A.Z-H2B.V nucleosomes are less able to stall MNase digestion than canonical nucleosomes (Figure 3), a difference that likely occurs due to reduced positive charges at SHL 3.5 as well as other changes to the histone-DNA interface. Reduced constraint on nucleosomal DNA may facilitate its displacement by RNA polymerase II, lowering the barrier to initiation and early elongation and reducing the energetic cost of repeated polymerase engagement. Thus, the pronounced instability of H2A.Z-H2B.V nucleosomes could directly contribute to the constitutively active transcriptional environment of *T. brucei*. In other eukaryotes, H2A.Z nucleosomes are actively removed by INO80-family chromatin remodellers (109,110), the chaperone ANP32E (111), and transcription-coupled turnover mechanisms (112). Trypanosomes possess a SWR1-related complex that deposits H2A.Z but no identified eviction machinery (26,28). One possibility is that intrinsic instability may lower the energy barrier to transcription sufficiently that a dedicated eviction pathway becomes dispensable.

Despite their instability, H2A.Z-H2B.V nucleosomes retain comparatively well-ordered entry/exit DNA relative to canonical trypanosome (19) and metazoan H2A.Z nucleosomes (7,31), indicating that core stability and DNA trajectory are separable properties. Flexible DNA ends can permit closer packing and stacking of adjacent nucleosomes (20,31,82). Such interactions are refractory to transcription and have been proposed to stabilise broad H2A.Z domains in repressive heterochromatin (31). Trypanosome H2A.Z domains are similarly broad, spanning ∼10 kb across polycistronic transcription units and into the first genes (14). We hypothesise that the ordered ends of *Tb* H2A.Z-H2B.V nucleosomes (Figure 4) and open nucleosome arrays (Figure 5) exert the opposite effect and remain permissive for constitutive transcription. At the same time, these nucleosomes may have a higher propensity for long-range, inter-nucleosome interactions based on their altered magnesium-dependent behaviour (Figure 5). *T. brucei* TSSs form spatially clustered nuclear domains (27) and the intrinsic biophysical properties of variant nucleosomes described here may contribute directly to that organisation.

The *Tb* H2A.Z-H2B.V acidic patch is likewise structurally distinct. Whereas H2A.Z nucleosomes in other eukaryotes carry a modestly expanded patch that modulates compaction and effector binding (31,84), the *Tb* H2A.Z-H2B.V acidic patch is both larger and has a more anionic electrostatic surface that differentially modulates enzyme activity (Figure 6). Interestingly, both KKT23 and DOT1B appear to be more active on H2A.Z-H2B.V nucleosomes (Figure 4 & 6D) although we cannot discount technical assay limitations that may affect this result (e.g. due to antibody specificity). While KKT23 target sites on H2A.Z have not been mapped, DOT1B-mediated H3K76 trimethylation is only modestly enriched at TSS (25,70). The enhanced activity of these enzymes could therefore serve to maintain histone marks against transcription-coupled turnover of the variants rather than to establish variant-specific modification patterns.

Additionally, the altered acidic patch could provide a mechanism for specific recruitment to TSSs. Our H2A.Z-H2B.V interactome (Figure 7) is dominated by factors previously mapped to trypanosome TSS (28,88,103), particularly bromodomain proteins and histone acetylation and deacetylation complexes. Acetylation neutralises positive charge, weakening histone-DNA contacts (113) and promotes chromatin decompaction (114,115). A combination of H2A.Z and histone PTMs have also been shown to enhance the activity and binding of effector proteins in model eukaryotes (116–118). Acetylation of the Tb H2A.Z N-terminal tail controls transcriptional output (25), but the chromatin structure and events that allow constitutive transcription remain unmapped. In this context, the strong enrichment of a putative SIN3-like HDAC1 complex (28) is initially counterintuitive given the acetylation-rich environment of TSSs (25). We suggest a model where dynamic cycles of acetylation and deacetylation could help activate and reset TSSs to allow constitutive transcription. Future work looking at H2A.Z-H2B.V histone marks at TSSs would be invaluable to understand whether histone acetylation plays a structural and/or signalling role in trypanosome transcription initiation.

Together, our results demonstrate that H2A.Z-H2B.V nucleosomes constitute a structurally and functionally distinct chromatin state. By integrating obligate histone pairing, reduced stability, an altered acidic patch, and selective recruitment of chromatin regulators, these nucleosomes define transcription start regions in *T. brucei*. More broadly, our findings highlight how histone variant evolution can rewire chromatin architecture to generate lineage-specific solutions for transcriptional control in the absence of canonical promoter-based regulation.

## Data Availability

The cryo-EM density map and associated meta data for the *T. brucei* H2A.Z-H2B.V NCP have been deposited at the Electron Microscopy Data Bank under accession number EMD-57961. Raw micrographs have been uploaded to EMPIAR-13592. The atomic coordinates of the structure of the *Tb* H2A.Z-H2B.V NCP have been deposited in the Protein Data Bank under accession number 30QQ. Mass spectrometry proteomic data has been uploaded to PRIDE database under identifier PXD084419.

## Acknowledgements

We thank Bungo Akiyoshi and members of the Wilson lab for helpful discussions and critical reading of the manuscript. We thank David Kelly for help with the fluorescence microscopy imaging. This work was also supported by the Edinburgh Protein Production Facility (EPPF). We are grateful to Thomas Schalch for the MMTV and 12×177bp Widom 601 plasmid. Thanks to Iain Manfield, Centre for Biomolecular Interactions, University of Leeds for access to FIDA bio instrumentation and Thomas Bedwell and Michael Singh for optimisation and advice on FIDA Bio experiments. We are very grateful to Peter Harrison and Diamond Light Source for access and support of the Cryo-EM facilities at the UK national electron bio-imaging centre (eBIC), proposal EM-BI31827, funded by the Wellcome Trust, MRC and BBSRC. We thank Hannah Wapenaar for human H2A.Z.1 octamers and Christina Cardenal Peralta for help with mass spectrometry samples. We are grateful to Roberta Carloni, Robin Allshire and Keith Matthews for providing procyclic trypanosome cells and Patryk Ludzia and Bungo Akiyoshi for the KKT23 protein.

## Author contributions

MDW conceived the study and supervised the project. GDÜ and MDW designed the experiments, analysed the data, and wrote the manuscript, with input from the other authors. Unless otherwise stated, GDÜ purified the DNA and protein components and performed all biochemistry experiments. MDW performed the FIDA Bio experiments. Cryo-EM data collection and image processing was performed by MDW and GDÜ. Model building was performed by GDÜ. Initial cryo-EM screening was performed by MS. CS performed mass spectrometry sample loading and raw data analysis.

## Supplementary Figure Legends

**Supplementary Figure 1: Histone H2A.Z and H2B.V divergence in eukaryotes**

**A.** Pairwise protein sequence identity matrix of histone H2A and H2A.Z variants in various eukaryotes.

**B.** Pairwise protein sequence identity matrix of different eukaryotic histone H2B variants. Canonical histones are labelled with a grey “C”.

**Supplementary Figure 2: Conservation of histones H2A.Z and H2B.V in kinetoplastids**

Multiple sequence alignments of kinetoplastid H2A.Z (**A**) and H2B.V (**B**). The secondary structure of *T. brucei* histones is shown above each alignment and was defined based on the structure of the *T. brucei* H2A.Z-H2B.V NCP. Residues corresponding to alpha helices are shown in boxes. Specific residues involved in nucleosome stability or interactions are category highlighted (top legend) and numbered based on *T. brucei* H2A.Z and H2B.V sequences. Alignment gaps are shown in grey.

**Supplementary Figure 3: *In vitro* reconstitution of H2A.Z-H2B.V NCPs and their overall stability**

**A-C.** Size exclusion chromatography (SEC) traces (left) and SDS PAGE analysis of fractions from each SEC trace (right) of H2A.Z-H2B.V octamer reconstitution attempts. **A.** Combining His_6_-H2A.Z, H2B.V, H3, and H4 produced multiple species (peaks a, b and c). Reconstitution of H3-H4 tetramers (**B.**) and His_6_-H2A.Z, H2B.V dimers (**C.**) was used to confirm the identity of peaks in panel A.

**A. D.** An overlay of the SEC traces shown in panels A-C.

**B. E.** Native (left) and SDS PAGE (right) gels of His_6_-H2A.Z-H2B.V NCPs reconstituted with octamers or a mixture of dimers and tetramers.

**C. F.** Representative native (left) and SDS (right) PAGE gels of NCPs reconstituted with Widom 601 145 bp DNA that were used for TDA assays.

**D. G.** Comparison of melting temperature (T_m_) values obtained for different *T. brucei* and human NCPs (n = 3). Data for human canonical NCPs was taken from our previous publication (19) and is shown using hollow circles. The mean T_m_ value for each NCP is indicated with a black line.

**E. H.** Thermal denaturation curves of different NCPs (n = 3). RFU = relative fluorescence intensity. Error bars represent the standard error of mean.

**Supplementary Figure 4: Single particle cryo-EM processing pipeline for the H2A.Z-H2B.V NCP**

An overview of the cryo-EM processing pipeline for the H2A.Z-H2B.V NCP including particle picking, 2D classification, 3D classification, and subsequent refinement steps. Viewing direction distribution plots showing the representation of different particle orientations before and after particle rebalancing are shown. Orientation diagnostics metrics including the sampling compensation factor (SCF) and the conical FSC area ratio (cFAR) are also provided. The GS-FSC resolution and B-factor sharpening of the final model are stated at the bottom.

**Supplementary Figure 5: Cryo-EM reconstruction and model building of the H2A.Z-H2B.V NCP**

**A.** Representative cryo-EM micrograph of His_6_-H2A.Z-H2B.V NCPs.

**B.** Gold standard Fourier shell correlation (GSFSC) curve (left) and 3D-FSC curve (right) of the final cryo-EM reconstruction of His_6_-H2A.Z-H2B.V NCPs from CryoSPARC (49). In both curves, a horizontal black line depicts the threshold at FSC = 0.143. In the 3D-FSC curve, the green bars represent the relative frequency of 0.143 FSC crossings at different resolutions.

**C.** Cryo-EM density maps coloured by local resolution (colour scale indicated below) of the His_6_-H2A.Z-H2B.V NCP estimated using Local Resolution Estimation in CryoSPARC.

**D.** The model of the H2A.Z-H2B.V NCP fit into the final cryo-EM density map (left) and close-up views of model building for each histone and DNA (right). The chain and residue limits of each fragment are provided below.

**Supplementary Figure 6: Analysis of different histone-histone interactions in the H2A.Z-H2B.V NCP**

**A.** Magnified view of H2B(V)-H4 interactions at the dimer-tetramer interface of the canonical NCP (PDB: 8COM (19)) and H2A.Z-H2B.V NCP (this study).

**B.** Left: An overlay of DNA gyres from different NCP structures determined by single particle cryo-EM in previous studies. The structures include *Homo sapiens* (*“Hs”*) canonical NCPs (PDB: 7XD1 (80)), *Tb* canonical NCPs (PDB: 8COM (19)), *Hs* H2A.Z NCPs (PDB: 9B3P (119)), *Mus musculus* (*“Mm”*) H2A.Z NCPs (PDB: 7M1X (31)), *Arabidopsis thaliana* (*“At”*) H2A.Z NCPs (PDB: 9K43 (11)), and *Tb* H2A.Z-H2B.V NCPs (this study). The structure of the H2A.Z-H2B.V octamer is included for reference and shown with surface representation. The distinctive cavity near SHL 5 is indicated with a dashed oval. *Right*, Distance measurements of NCP inter-gyre distance at SHL 4.5 and 5.5 are shown as dashed orange lines. Four pairwise distances were measured between the phosphate atoms of nucleotides I:34, I:35 and J:45, J:46 (SHL 4.5) as well as nucleotides I:24, I:25 and J:56, J:57 (SHL 5.5). “I” and “J” indicate DNA chains. For PDB 9K43, chains I and J were swapped to measure equivalent positions. Distances are plotted on the left, equivalent measurements are connected by dashed lines and median values of distance across 4 measurements highlighted by a horizontal line.

**C.** Size exclusion chromatography profiles of histone dimers reconstituted with different combinations of canonical and variant histones. Solid lines = canonical-canonical or variant-variant combinations; dashed lines = canonical-variant combinations.

**D.** Dimer formation propensity of each histone combination (n = 2). The y-axis represents the ratio of the maximum A280 value of a dimer peak and the maximum A280 value of a soluble aggregate peak from two independent SEC traces obtained for each histone combination. For H2A.Z-H2B.V, one data point (hollow circle) is from dimers reconstituted with tagged His_6_-H2A.Z and H2B.V.

**E-F.** Predicted interfaces between H2A-H2B.V (A) and H2A.Z-H2B (B) compared to experimentally determined H2A.Z-H2B.V interfaces. The predicted interfaces were generated by overlaying *Tb* canonical and H2A.Z-H2B.V structures. Key residues are labelled. Potential clashes are shown with dashed lines. Canonical NCP = PDB: 8COM (19), H2A.Z-H2B.V NCP = this study.

**Supplementary Figure 7: *In vitro* reconstitution and quality control of 12-mer nucleosome arrays**

**A.** Schematic representation of different nucleosome arrays reconstituted with 12x Widom 601 177 bp DNA (black line), histones (grey circle = any octamer, red circle = *T. brucei* canonical octamer), and fluorophores (blue star = Alexa647, green star = OGG488). Fluorescent nucleosome arrays were reconstituted by combining unlabelled and fluorescently labelled octamers at a 1:12 ratio (middle) or by using fluorescently labelled octamers alone (right).

**B.** Representative native agarose gels of various nucleosome arrays reconstituted using the methods shown in panel A. 147 bp MMTV competitor DNA was added to promote array saturation and MgCl_2_ was added to precipitate saturated arrays and thereby remove free DNA. The arrays are shown before (input) and after (pellet) magnesium precipitation cleanup.

**C.** AccI restriction enzyme digestion of nucleosome arrays. The position of AccI recognition sites in the histone-bound Widom 601 repeats is shown in the schematic on top. Agarose gels showing array DNA and deproteinated nucleosome arrays in the absence (-) and presence (+) of AccI are shown below.

**D.** ScaI restriction enzyme digestion of nucleosome arrays. The position of ScaI recognition sites in the linker DNA is shown in the schematic on top. Native PAGE of array DNA and nucleosome arrays in the absence (-) and presence (+) of ScaI are shown below. In each gel, the section containing 177 bp ScaI digestion products is outlined with dashed lines and shown underneath in a blue box.

**Supplementary Figure 8: Biophysical characterisation of H2A.Z-H2B.V nucleosome arrays**

**A-B**. Representative raw (left) and fit (right) Taylor dispersion trace from FIDA Bio experiment for canonical OG488 (A) and H2A.Z nucleosome arrays (B) with a polydispersity index (PDI) and regression model fit to the data (R^2^) given.

**C.** Fluorescence microscopy of 1/12 Alexa647-labelled nucleosome arrays with and without 2.5 mM magnesium chloride treatment prior to imaging.

**Supplementary Figure 9: The pipeline and dataset characteristics of the AP-MS approach used to identify nucleosome interactors in *T. brucei***

**A.** The number of unique proteins identified in each condition and replicate. Due to a low number of proteins, replicate 1 from the Canonical+GA sample was omitted from further analyses.

**B.** Schematic representation of the AP-MS pipeline showing the number of unique proteins identified across all conditions and replicates after different stages of filtering. The “≥ 2 peptides” filter selected for proteins identified with at least two or more unique peptides. After this stage, replicate 1 from the Canonical+GA was omitted. The “DIA Analyst” filter removed proteins with a high number of missing values using the automatic pre-filtering step in the DIA analyst software (Monash Proteomics). The “Nuclear (TrypTag)” filter selected for proteins assigned to the GO term nucleus (GO:0005634) in the TrypTag database (91). The “Contaminants out” filter removed proteins that were described with the terms “ribosome”, “ribosomal”, “endoplasmic”, “endocytic”, “pore”, “mitochondrial”, “mitochondrion”, and/or “kinetoplast” in the TriTrypDB database (35). The “Nuclear (TrypTag)” and “Contaminants out” filters were only used for case-specific, qualitative analysis of the data. All other analyses were performed on data from the “DIA Analyst” filter (n = 3,048).

**C.** A correlation matrix showing Pearson’s correlation coefficients between mass spectrometry protein intensity values obtained for each condition and replicate post-imputation.

**D.** A violin plot showing the distributions of protein intensity values (Log_10_ scale) obtained for each condition and replicate post imputation. A black line representing median intensity is drawn through each dataset.

**Supplementary Figure 10: Key nucleosome interactors identified across the AP-MS dataset**

Volcano plots showing pairwise comparisons of different conditions (Canonical, Canonical+GA, H2A.Z-H2B.V, and H2A.Z-H2B.V+GA). The missing comparison between Canonical and H2A.Z-H2B.V samples is shown in Figure 7B. In each plot, consistent H2A.Z-H2B.V binders are shown in yellow and Canonical binders are shown in red. Relevant histones and key interactors of Canonical and H2A.Z-H2B.V nucleosomes are highlighted if they appear above the P-value threshold of 0.05 (horizontal dashed line) and Log2 Fold Change threshold of |1| (vertical dashed lines).

**Supplementary Figure 11: Additional data related to the identification of key H2A.Z-H2B.V and canonical nucleosome interactors**

Box plots from DIA-Analyst (Monash Proteomics) showing measured (circles) and imputed (triangles) Log_2_-transformed intensities obtained for key interactors of H2A.Z-H2B.V nucleosomes (highlighted yellow) and canonical nucleosomes (highlighted pink). “Can” = Canonical, “ZBV” = H2A.Z-H2B.V. The y-axis range in each graph has been scaled differently based on the data.

**Supplementary Figure 12: Additional data related to the identification and analysis of HDAC1 complex subunits as H2A.Z-H2B.V interactors**

**A.** Box plots from DIA-Analyst (Monash Proteomics) showing measured (circles) and imputed (triangles) Log_2_-transformed intensities obtained for each member of the *T. brucei* HDAC1 complex across all conditions (“Can” = Canonical, “ZBV” = H2A.Z-H2B.V). Please note that the y-axis range in each graph has been scaled differently based on the data.

**B.** A structural alignment of the AlphaFold2-predicted structure of *Tb* Q57TZ5 (120) and experimentally determined structures of SIN3 proteins including human SIN3B (PDB: 8C60 (121), chain A) (121), the *Schizosaccharomyces pombe* SIN3 homolog Pst1 (PDB: 8I03 (122), chain A) (122), and *Saccharomyces cerevisiae* SIN3 (PDBs: 8TOF, chain A and 8GA8, chain A) (123,124) performed using TM-Align on the RCSB server (42).

**C.** The AlphaFold2-predicted structure of *Tb* Q57TZ5 coloured by the predicted local distance difference test (pLDDT) score (120).

**D.** The AlphaFold2-predicted structure of *Tb* Q57TZ5 (120) aligned to experimentally determined structures of human and yeast SIN3 proteins from the structural alignment in panel B.

**Supplementary Figure 13: Histone acetylation in the AP-MS dataset**

**A.** Top: Percentage of peptide acetylation identified at different histone modification sites. Each datapoint represents a single acetylated peptide and has been normalised by the total intensity of peptides obtained for the histone of interest in the repeat where the acetylated peptide was identified. Hollow data points represent peptides found in one of the Canonical or Canonical+GA repeats and fully coloured data points represent peptides found in one of the H2A.Z-H2B.V or H2A.Z-H2B.V+GA repeats. Bottom: A table summarising the repeats in which acetylated peptides were identified (grey boxes).

**Supplementary Table S1:** Cryo-EM data collection, processing, and model building parameters.

|  |  |
| --- | --- |
| **Data Collection** |  |
| Microscope | TFS Titan Krios |
| Detector | Gatan-K3 |
| Acceleration voltage (kV) | 300 |
| Number of micrographs | 13,861 |
| Frames per micrographs | 48 |
| Exposure time (s) | 2.3 |
| Dose per frame (e- Å<sup>-2</sup>) | 1.03 |
| Accumulated dose (e- Å<sup>-2</sup>) | 49.53 |
| Defocus range (µm) | -2.75 to -1.25 (0.25 steps) |
| **Frames** |  |
| Alignment software | Patch Motion Correction (CryoSPARC) |
| Frames used in final reconstruction | 0-48 |
| Dose weighting | Empirical (from Reference-Based Motion Correction, CryoSPARC) |
| **Contrast Transfer Function (CTF)** |  |
| CTF Fitting software | Patch CTF (CryoSPARC) |
| CTF Correction | Local (optimisation of per-particle defocus) |
| **Particles** |  |
| Picking software | Template-based picking (CryoSPARC) |
| Particles picked | 3,094,412 |
| Particles used in final reconstruction | 277,951 |
| **Alignment** |  |
| Alignment software | CryoSPARC |
| Initial reference map | Ab initio reconstruction |
| **Reconstruction** |  |
| Reconstruction software | CryoSPARC |
| Box Size (pixels) | 304 |
| Voxel size (Å) | 0.829 |
| Symmetry | C1 |
| Resolution estimate (Å) | 2.77 (GS-FSC) |
| Masking | Yes |
| B-Factor Sharpening (Å<sup>2</sup>) | -80 |
| cFAR | 0.64 |
| SCF | 0.85 |
| **EMD ID** | **57961** |
| Model Building |  |
| Number of protein residues | 755 |
| Number of DNA residues | 286 |
| Bond length outliers > 4σ (□) | 0 out of 12,584 |
| Bond length RMSD (□) | 0.008 |
| Bond angle outliers > 4σ ($^\circ$) | 0 out of 18,219 |
| Bond angle RMSD ($^\circ$) | 1.272 |
| Ramachandaran favoured \| allowed \| outliers (%) | 98.11 \| 1.89 \| 0.00 |
| Rama-Z (RMSD) whole \| helix \| loop | -0.17 (0.27) \| 0.21 (0.20) \| -0.67 (0.40) |
| Rotamer outliers (%) | 0.00 |
| CaBLAM outliers (%) | 0.28 |
| Clash Score | 1.94 |
| Model vs Data CC (mask) | 0.84 |
| EMringer | 4.29 |
| FSC Model vs Map 0.5 (unmasked) | 2.9 |
| MolProbity Score | 0.96 |
| **PDB ID** | **30QQ** |

