## Supplementary Figures for "H2A.Z-H2B.V nucleosomes form a specialised scaffold for transcription initiation in *Trypanosoma brucei*"

### Supplementary Figure 1

A

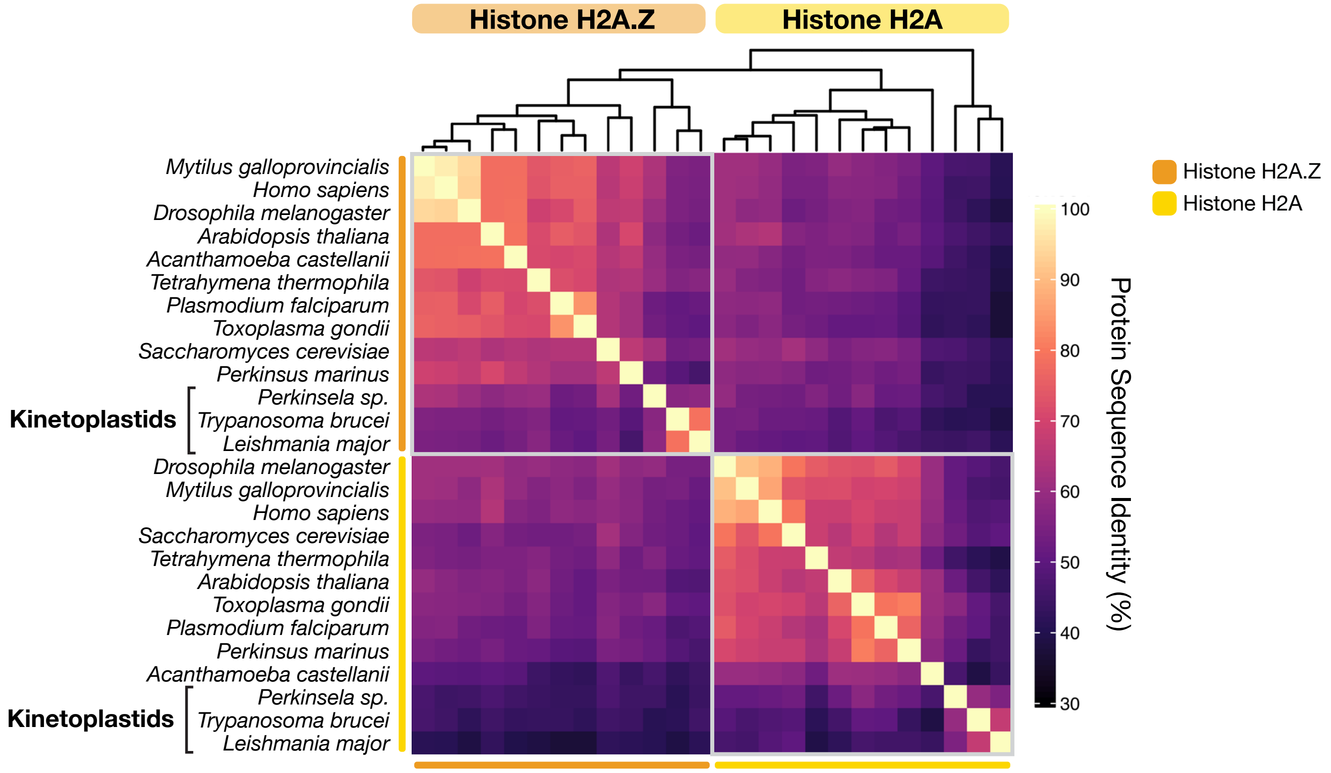


B

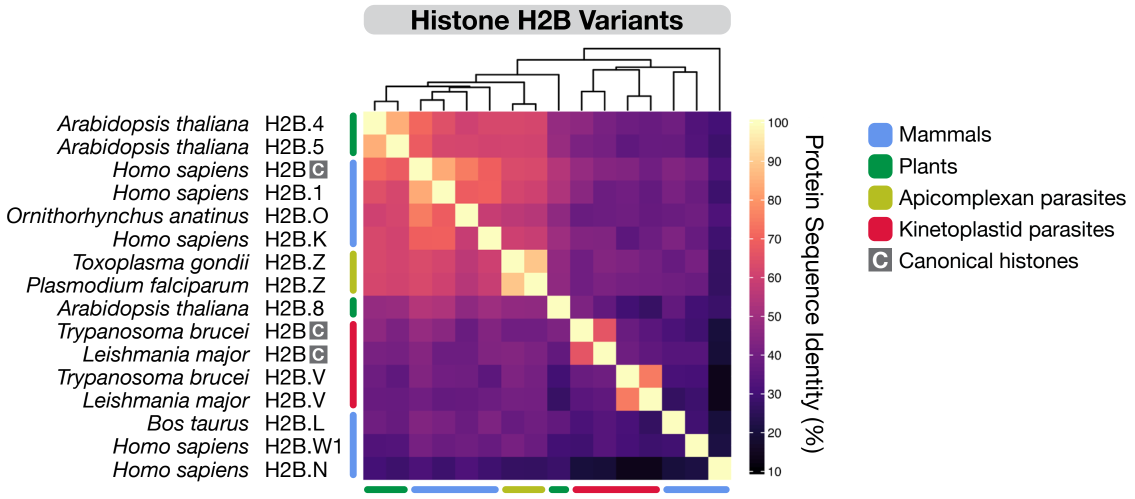

### Supplementary Figure 2

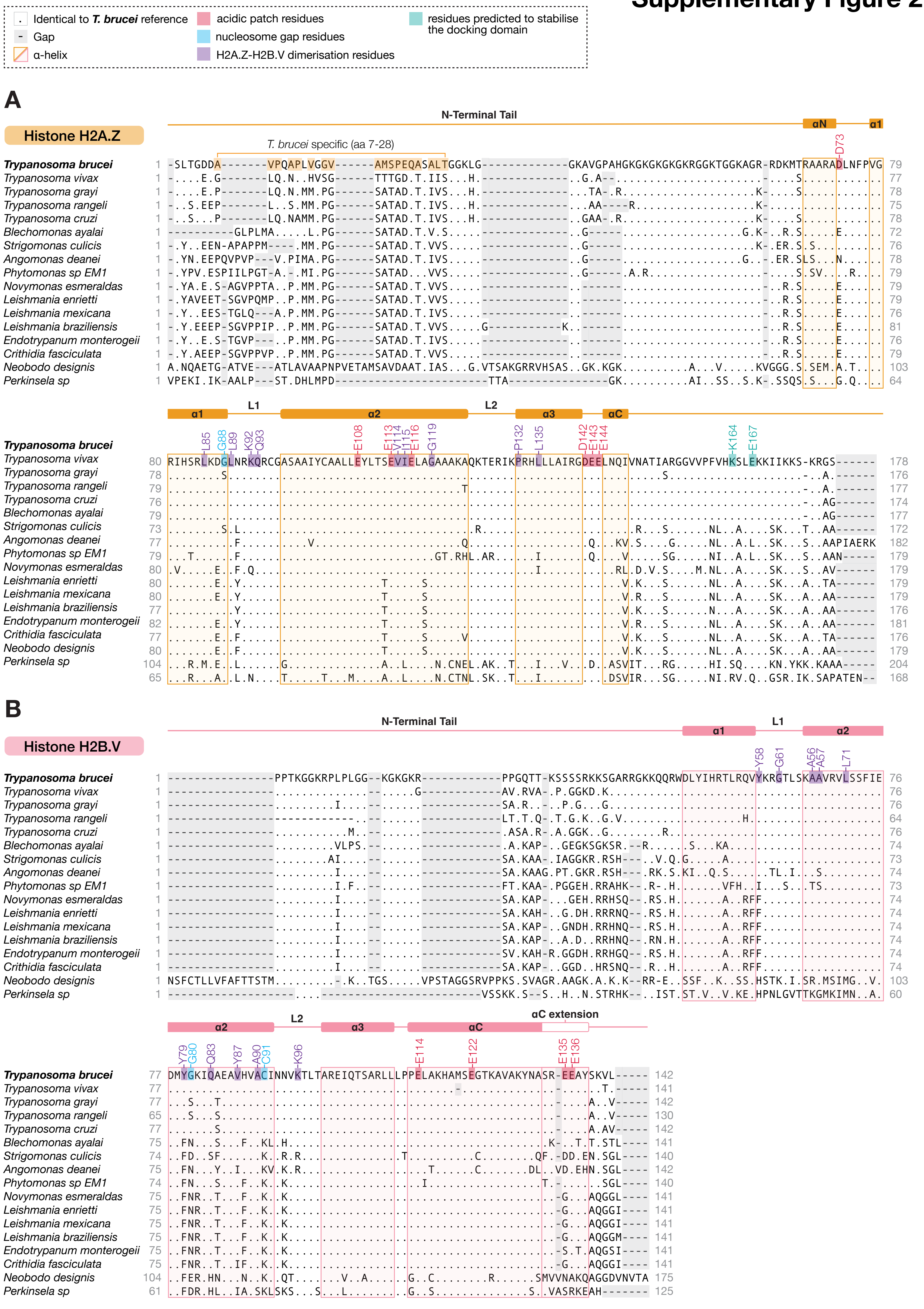

### Supplementary Figure 3

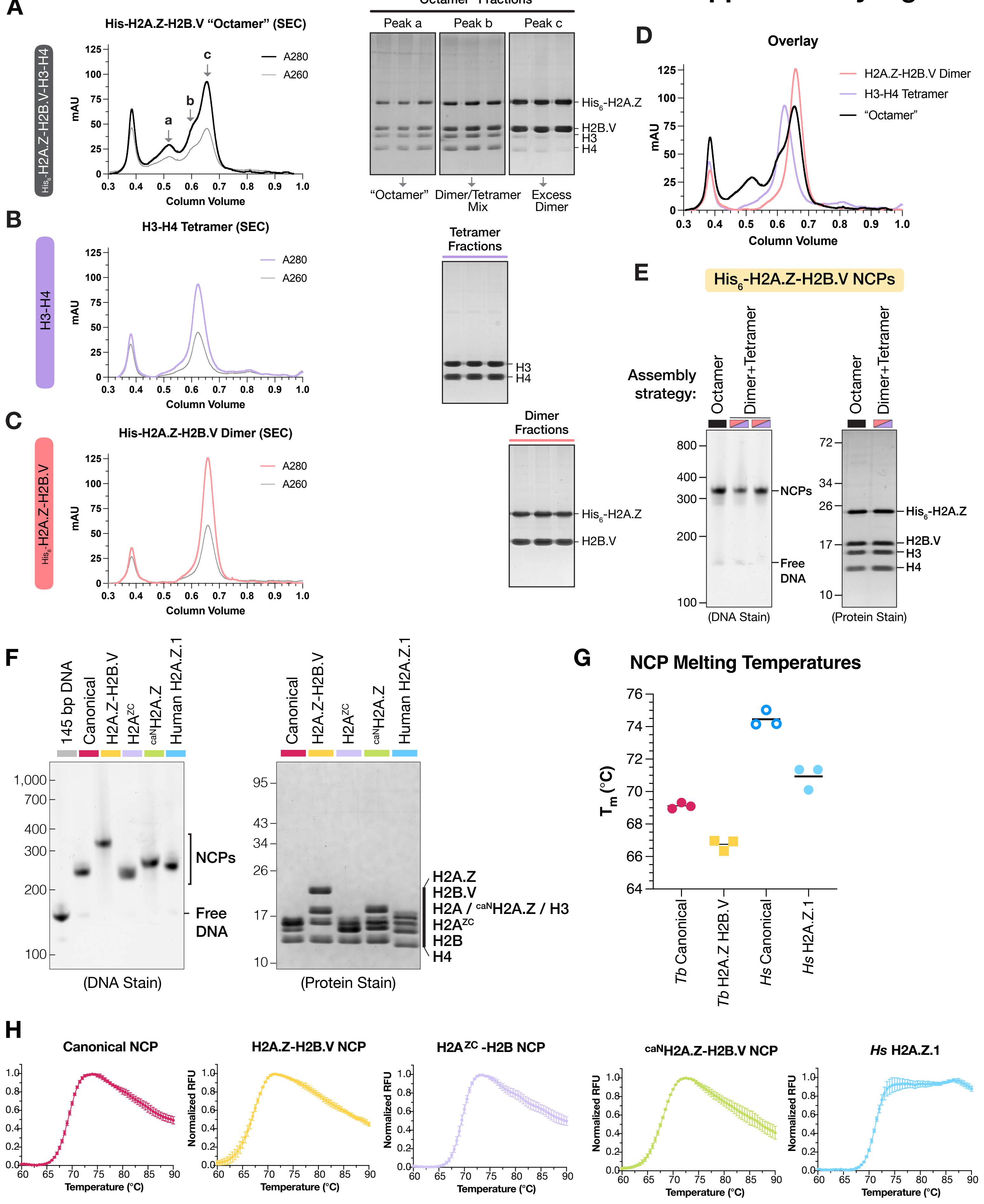

### Cryo-EM processing of the *T. brucei* H2A.V-H2B.V Nucleosome Core Particle

#### Supplementary Figure 4

**13,861 Movies**

**Patch Motion Correction**
**Patch CTF Estimation**

**Exposure curation**
9,924 Micrographs

**Micrograph denoiser**

**Manual Picking** (~1000 particles)
2D Classification
*templates*

**Template-Based Picking**
3,094,412 particles

**Particle Extraction**
256 px boxsize, bin 4

**2D Classification**
Round 1: 100 classes, 19 selected, 1,200,200 particles (43%)
Round 2: 100 classes, 48 selected, 1,055,641 particles (88%)

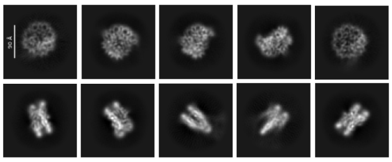

**Particle Extraction**
256 px boxsize

***Ab Initio Reconstruction***
Homogenous refinement
*starting model*

**Non-uniform Refinement 1**

**3D Classification 1**

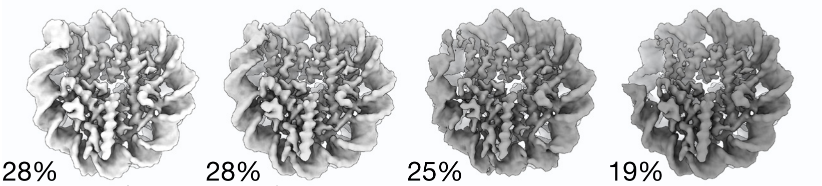


**Particle Extraction**
304 px boxsize, 574,920 particles

**Homogeneous Refinement 1**

**Reference-based Motion correction**

**Homogeneous Refinement 2**
*CTF refinement*

**3D Classification 2**

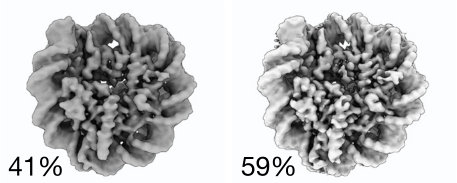


**Homogeneous Refinement 3**
*Optimise per-particle defocus*

**Orientation Diagnostics**
**& Rebalance Orientation**

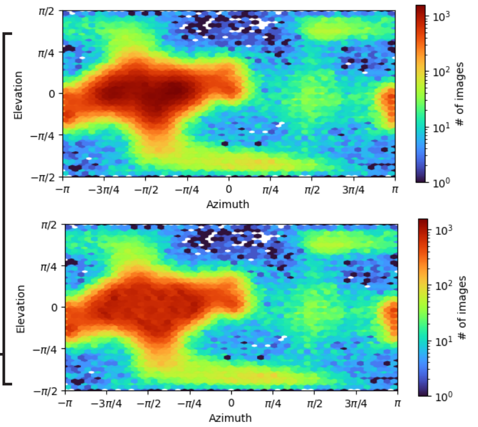


**Before Re-balancing**
cFAR = 0.62
SCF* = 0.78
339,596 particles

**After Re-balancing**
cFAR = 0.64
SCF* = 0.85
277,951 particles

**Homogeneous Refinement 4**

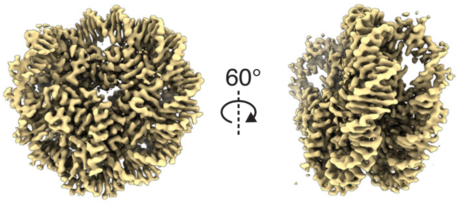


GS-FSC Resolution: 2.77 Å
B-Factor Sharpening: -80

### Supplementary Figure 5

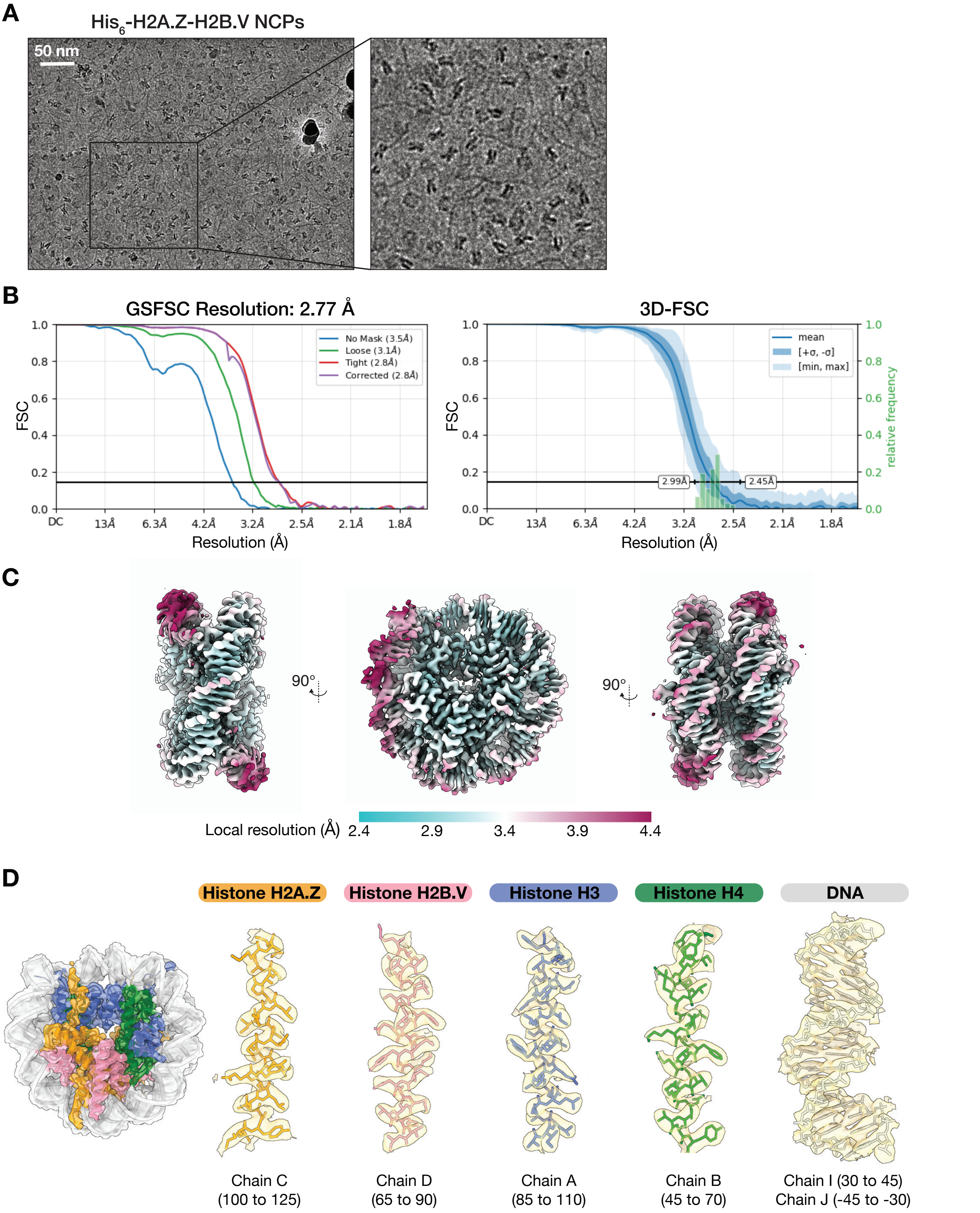

#### Supplementary Figure 6

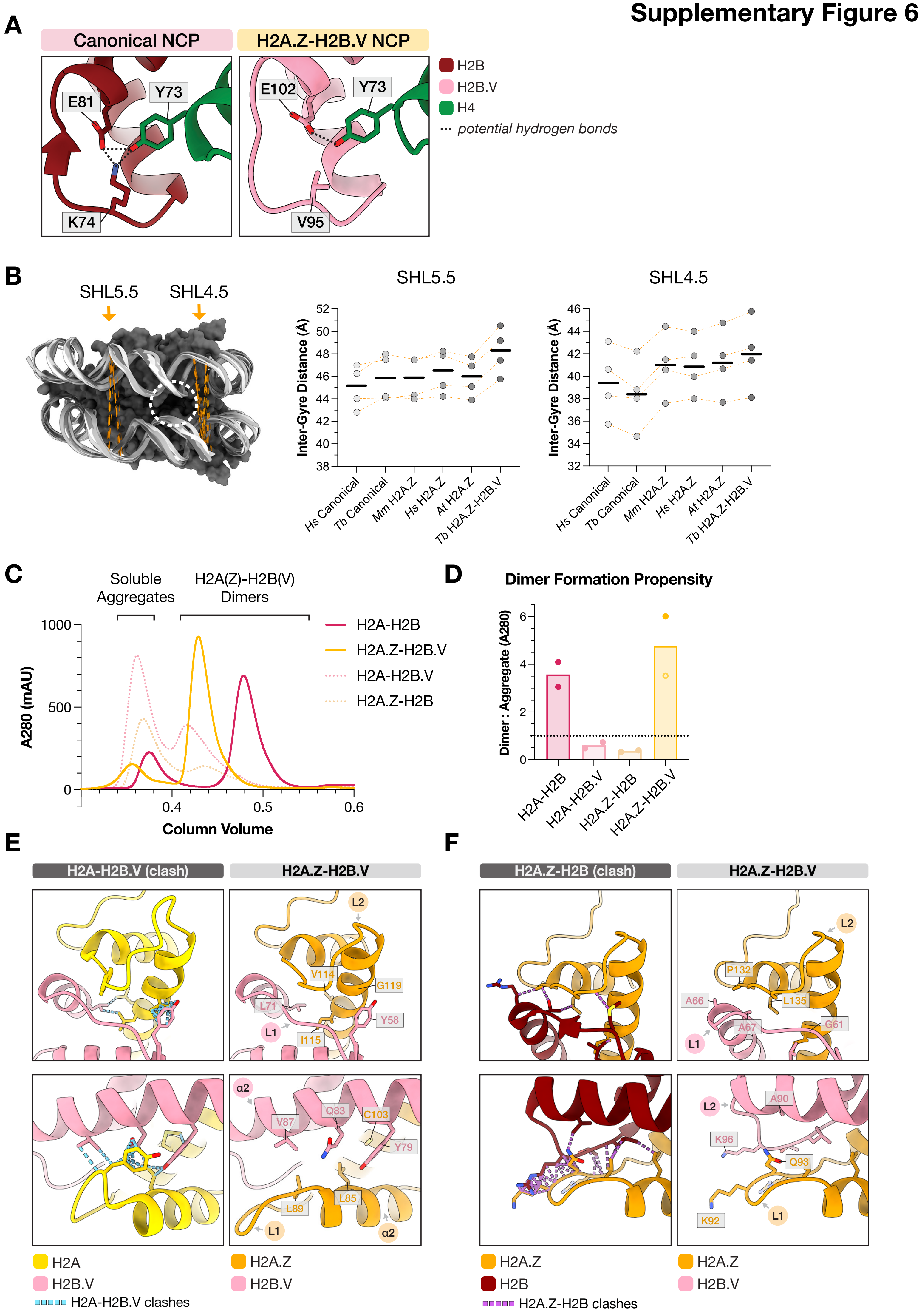

### Supplementary Figure 7

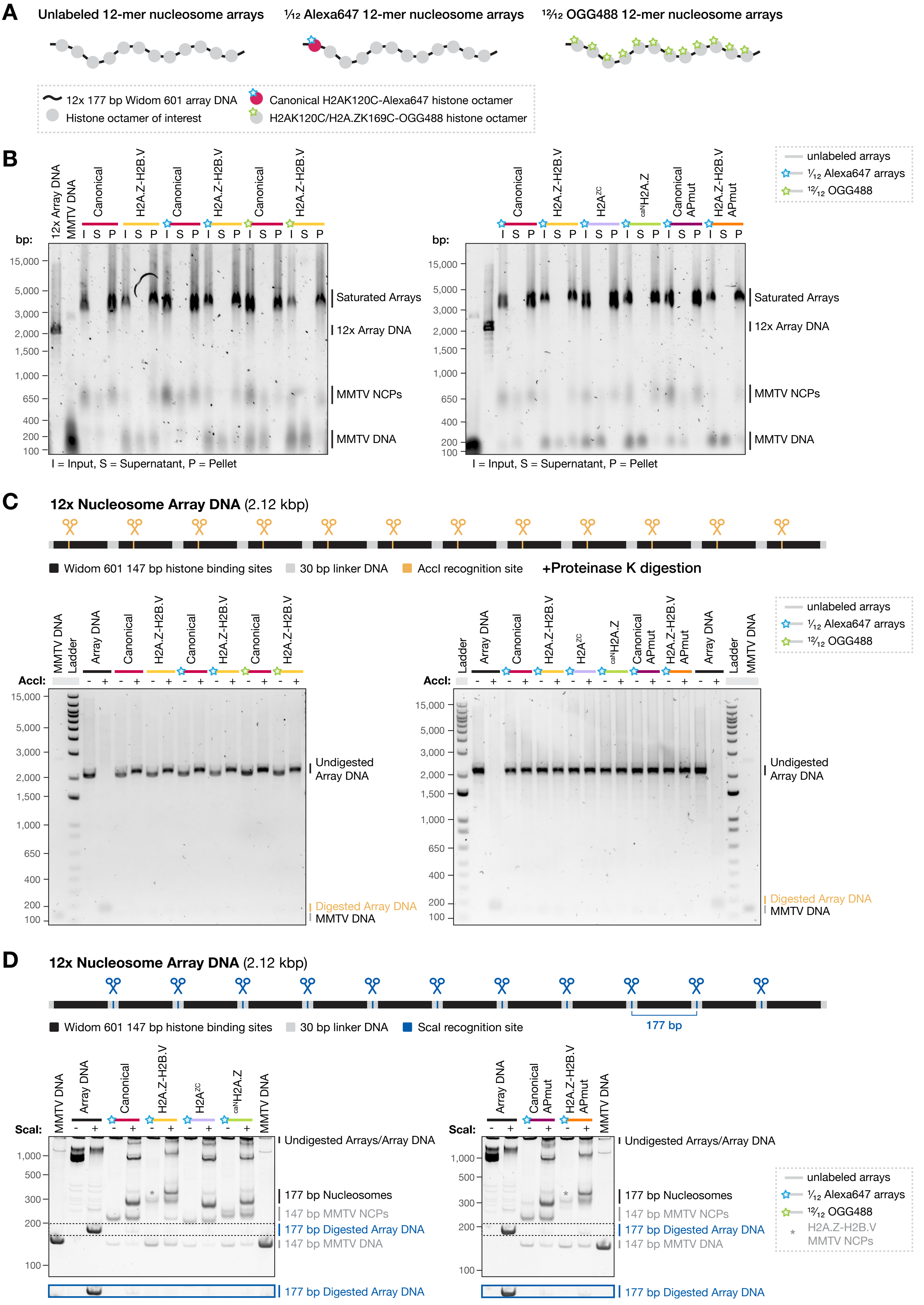

### Supplementary Figure 8

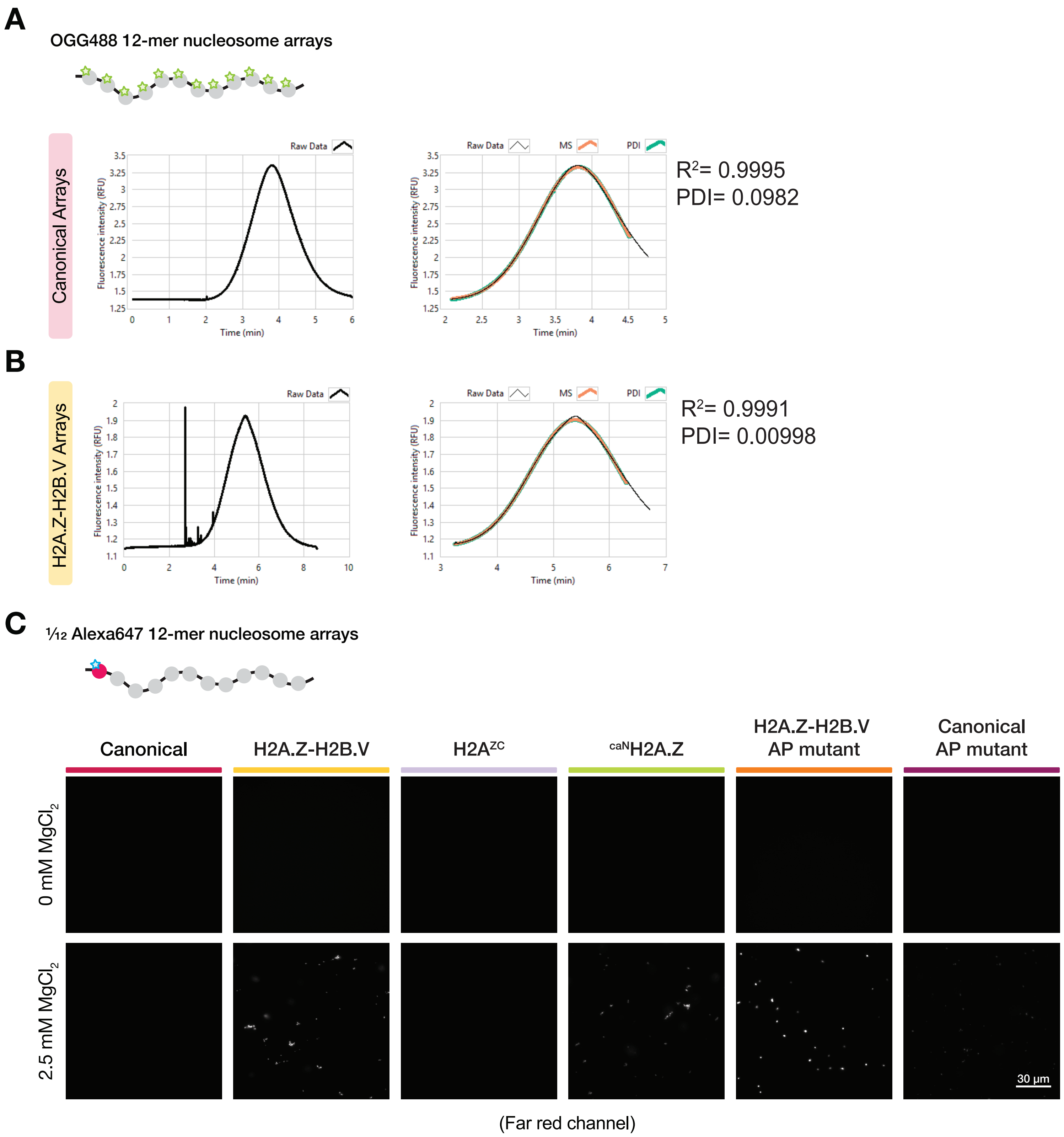

### Supplementary Figure 9

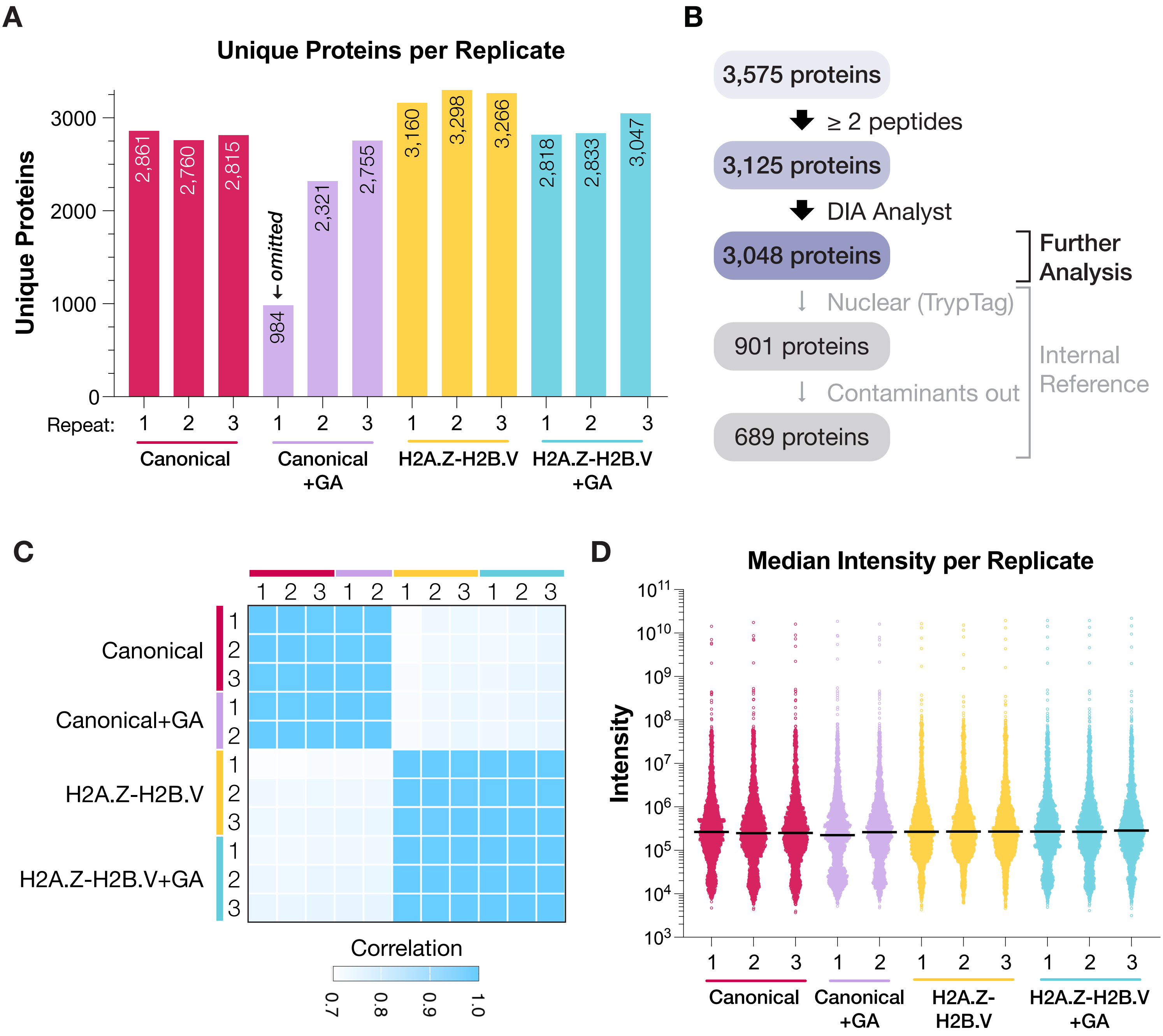

### Supplementary Figure 10

**A**

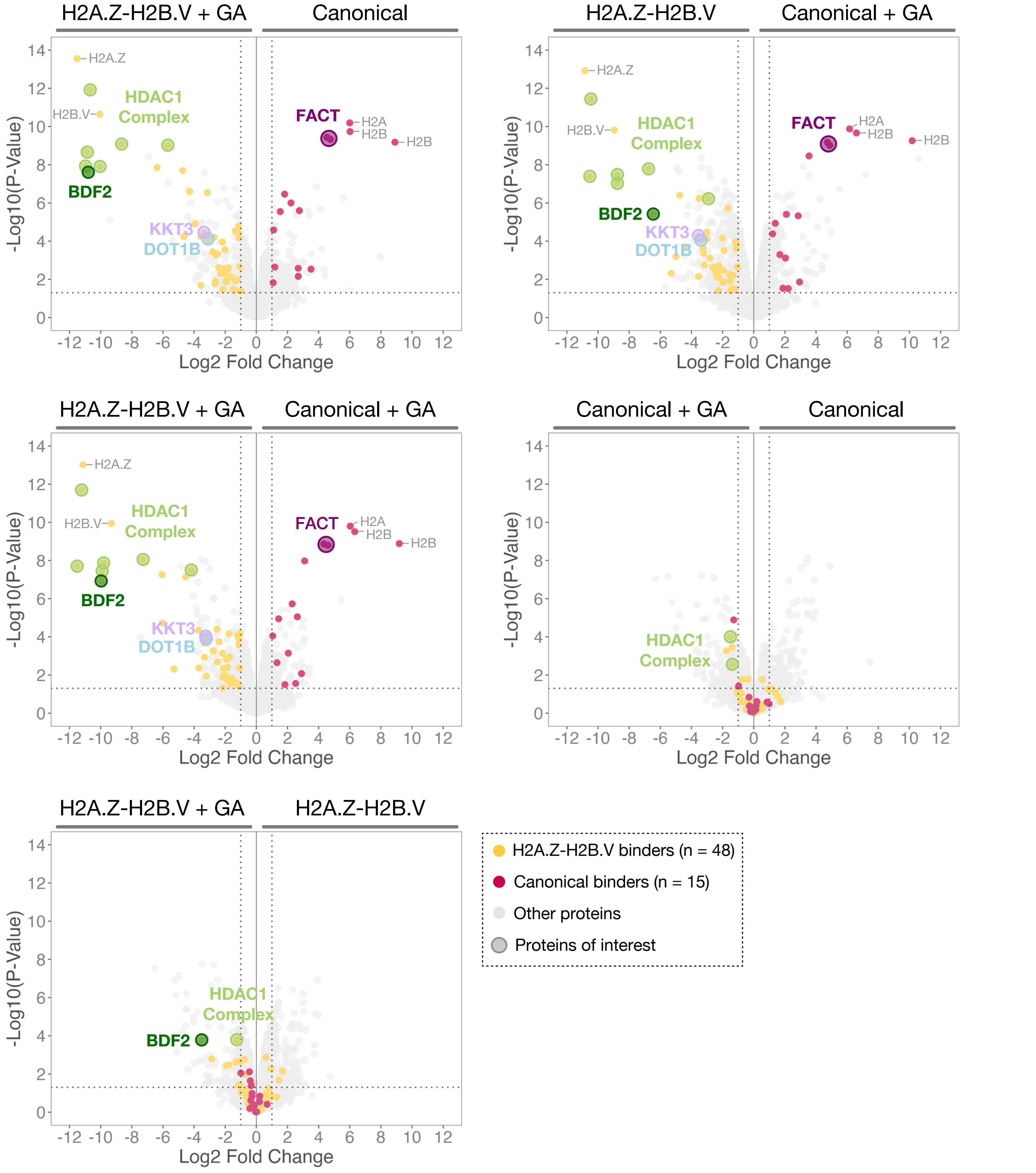

#### Supplementary Figure 11

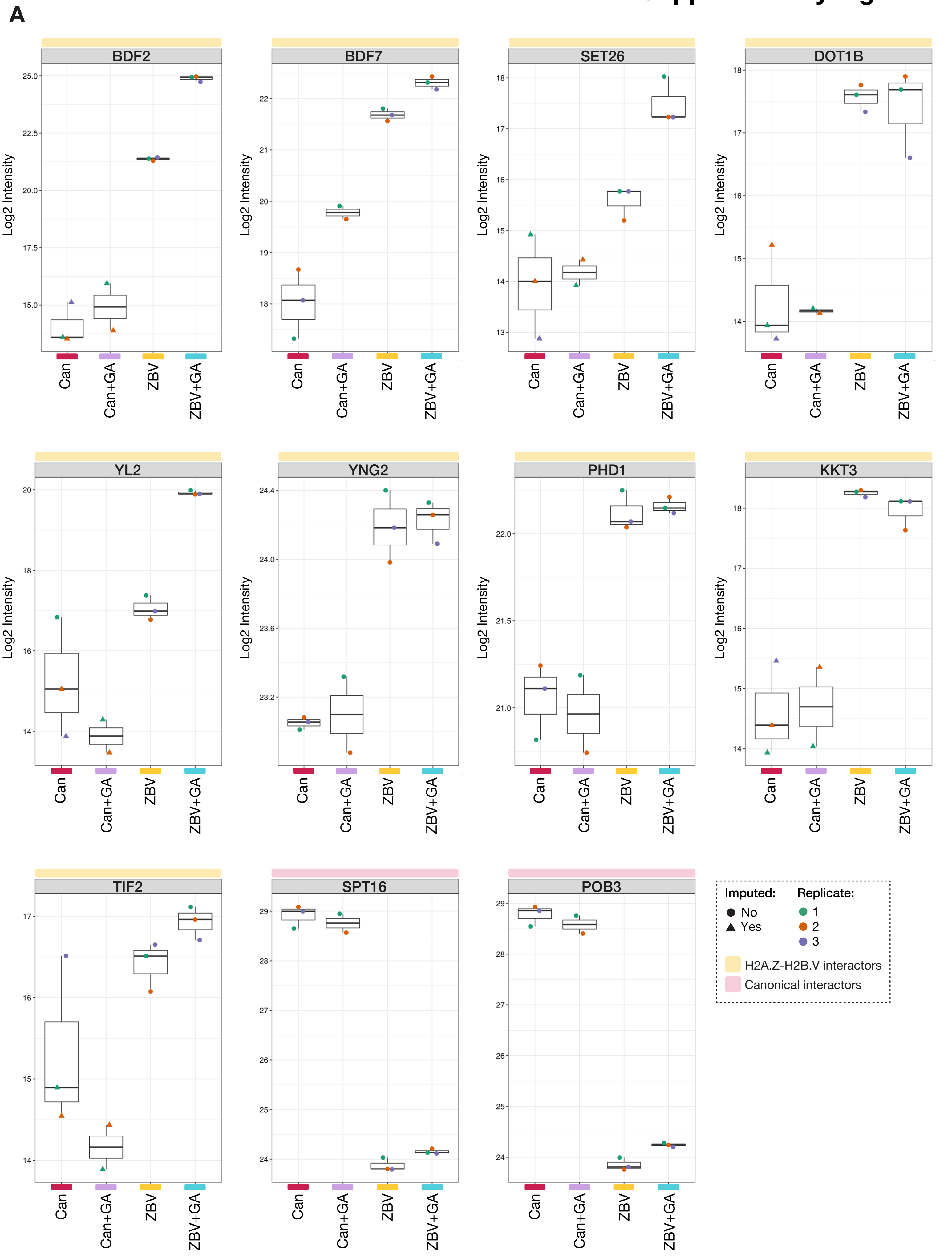

### Supplementary Figure 12

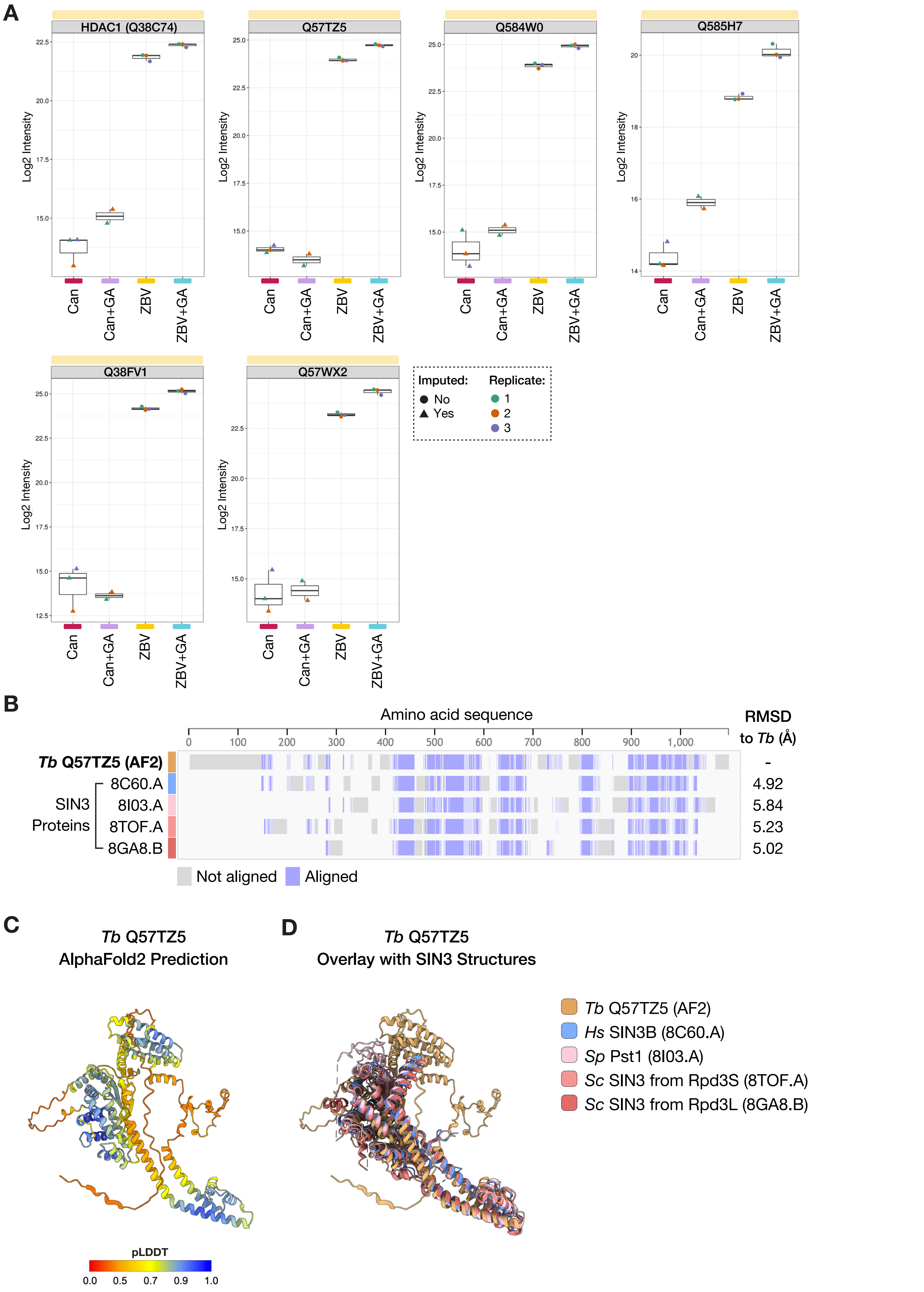

### Supplementary Figure 13

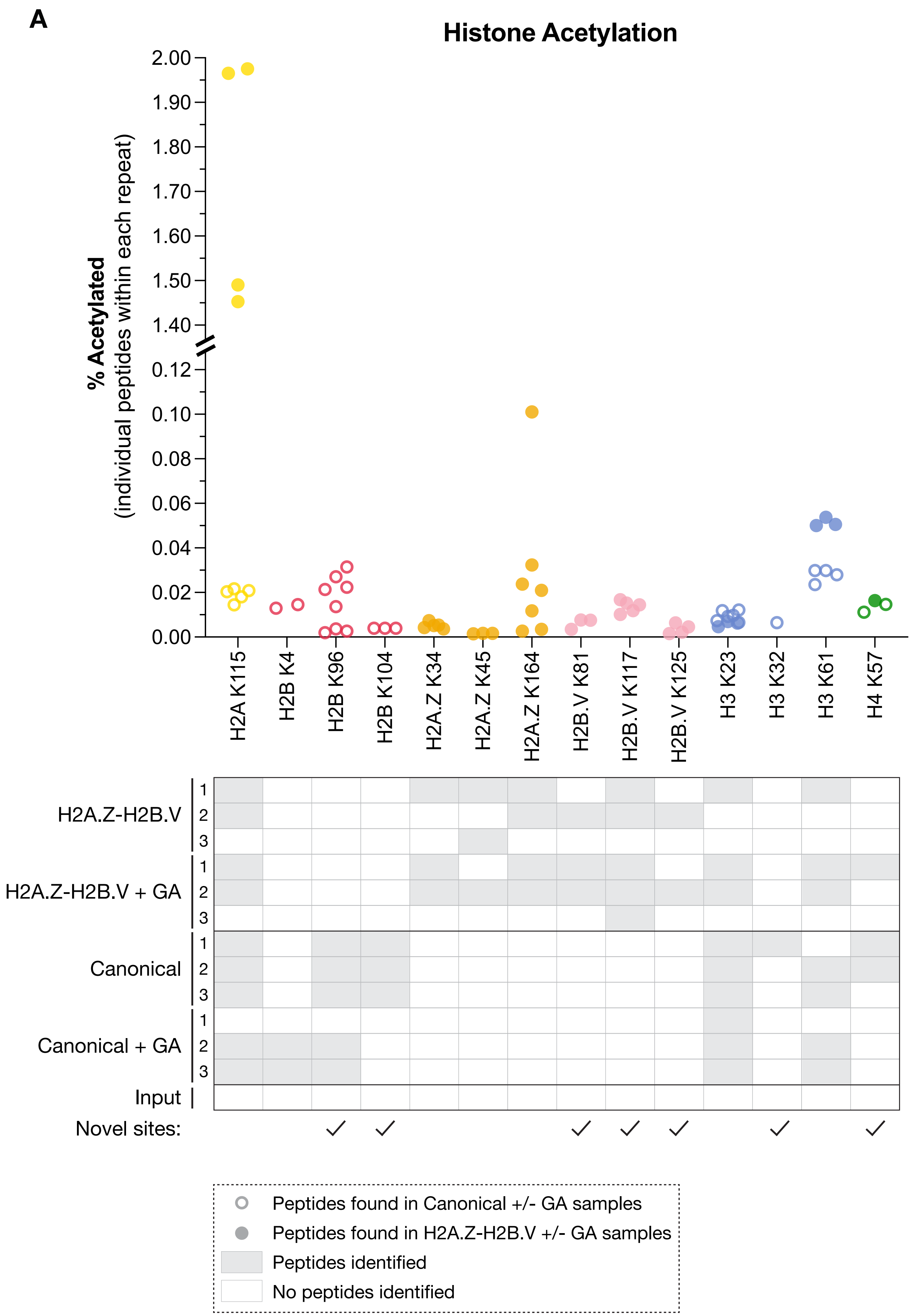
